# LIN28A-Dependent Kinome and Phosphoproteome Reprogramming Promotes Imatinib Resistance

**DOI:** 10.1101/2025.05.30.652226

**Authors:** Owen F.J. Hovey, Mallory I. Frederick, Quan M. Quach, Jenica H. Kakadia, Alyssa Wu, Kyle Yang, Tingting Wu, Xiang Ruan, Tomonori Kaneko, Courtney Voss, Ilka U. Heinemann, Shawn S. C. Li

**Affiliations:** Department of Biochemistry, Schulich School of Medicine and Dentistry, Western University, London, ON, N6A 5C1, Canada; Department of Oncology, Schulich School of Medicine and Dentistry, Western University, London, ON, N6A 5C1, Canada

## Abstract

Chronic myeloid leukemia (CML) resistance to BCR-ABL tyrosine kinase inhibitors (TKIs) can arise from ABL kinase domain mutations, BCR-ABL fusion gene amplification, or kinase-independent mechanisms. To investigate imatinib-resistance, we performed quantitative mass spectrometry comparing the proteome and phosphoproteome of K562 cells (a standard CML model) and ImR cells, an imatinib-resistant K562 derivative that also exhibits cross-resistance to second- and third-generation BCR-ABL TKIs. In addition to revealing global proteome and phosphoproteome changes associated with drug resistance, we identified LIN28A—a multi-functional RNA-binding protein—as a critical mediator of imatinib resistance. LIN28A was significantly overexpressed and hyperphosphorylated in ImR cells. Depleting LIN28A via shRNA restored imatinib sensitivity, while its ectopic expression in parental K562 cells induced imatinib resistance. Mechanistically, LIN28A coordinates an extensive kinase-substrate network regulating proliferation, survival, and metabolism to drive resistance. Notably, pharmacological inhibition of LIN28A-dependent kinases (PKC, AKT, SGK1, and RPS6K) suppressed ImR proliferation. Midostaurin, a clinical PKC/FLT3 inhibitor used in FLT3-ITD—positive AML, potently re-sensitized ImR cells to imatinib. Our findings suggest that targeting LIN28A and its downstream effectors, particularly PKC, could overcome resistance to imatinib and next-generation BCR-ABL inhibitors.

## INTRODUCTION

CML is a slowly progressive myeloproliferative neoplasm that accounts for approximately 15% of adult leukemia cases. In the United States, an estimated 8,860 new cases were diagnosis in 2022. Notably, the prevalence of CML has risen dramatically from ∼30,000 in 2000 to ∼150,000 in 2022, reflecting improved survival due to advances in therapy^1^. CML is characterized by a cytogenetic aberration resulting from a reciprocal translocation between chromosomes 22 and 9 (t(9;22)(q34;q11))^2^. This genetic rearrangement fuses the *breakpoint cluster region (BCR)* gene on chromosome 22 with the *ABL1* tyrosine kinase gene on chromosome 9, forming the oncogenic Philadelphia chromosome^2^. The resulting BCR-ABL1 oncoprotein exhibits constitutively tyrosine kinase activity due to loss of ABL1’s autoinhibitory domain. Through aberrant activation of proliferative and anti-apoptotic signaling pathways, the BCR-ABL1 fusion protein promotes uncontrolled myeloid cell expansion and survival, underpinning CML pathogenesis^3^.

Imatinib, the first tyrosine kinase inhibitor (TKI) approved by the US FDA for cancer therapy, revolutionized CML treatment by specifically targeting the oncogenic kinase BCR-ABL^4^. Indeed, the introduction of imatinib drastically reduced CML’s annual mortality rate from 10-20% to 1-2%^5^. Despite this remarkable success, only ∼10% of patients achieve treatment-free remission, leaving most dependent on lifelong TKI therapy^6^. Prolonged TKI therapy poses significant challenges, including accumulative toxicities and substantial financial costs and a proclivity for treatment resistance^1^. For instance, over 25% of patients require alternative TKIs due to intolerance or resistance, with 20-40% of resistant cases lacking clear mechanistic explainations^7^. Disease progression further complicates treatment efficacy—responsiveness to imatinib declines as patients transition from chronic phase to accelerated phase and blast crisis^8^. The most common resistance mechanism involves *ABL* kinase domain mutations (e.g., T315I), which has spurred the development of second- and third-generation TKIs such as dasatinib, nilotinib, bosutinib, and ponatinib^9^. However, clinical efficacy remains limited, with 10-40% of patients exhibiting refractoriness to these advanced therapies^10^. This persistent therapeutic challenge for CML highlights the critical need to elucidate alternative resistance mechanisms and develop next-generation treatment strategies.

While imatinib remains the first-line therapy for CML, resistance can arise through BCR-ABL-independent mechanisms^7^. These include upregulation of efflux pumps (e.g., ABC transporters) that actively remove imatinib from cells, activation of proliferative/pro-survival signaling cascades such as PI3K-AKT, MAPK, WNT, or JAK/STAT ^11–13^, and LIN28A-mediated pathways. The RNA-binding protein LIN28A, implicated in TKI resistance across multiple cancers, operate through let-7—dependent and/or let-7—independent mechanisms. The let-7 family of microRNAs are evolutionally conserved and play important roles in tumorigenesis^14^. In the let-7-dependent mode, LIN28A recruits TUT4/7 to polyuridylate and degrade let-7 miRNAs, resulting in upregulation of let-7 target genes^15^. Alternatively, LIN28 can bind directly to specific mRNAs and promote their transcription or degradation in a context-dependent manner^16–18^. The LIN28/let-7 axis contributes to therapeutic resistance in breast, lung, ovary, liver, and pancreatic cancers^16^. Notably, LIN28A is detected in 40% of advanced CML cases (accelerated phase/blast crisis) versus only 9% in chronic phase, suggesting its role in disease progression^19^.

Although imatinib has transformed CML treatment, approximately 10% of patients develop resistance^6^. The emergence of resistance to imatinib and subsequent generations of BCR-ABL TKIs underscores critical gaps in our understanding of resistance mechanisms. To address this deficiency, we established an imatinib-resistant K562 cell line (derived from a CML blast crisis patient) that exhibits cross-resistance to second- and third-generation inhibitors dasatinib and ponatinib. Comparative proteomic and phosphoproteomic analysis of parental versus resistant cells revealed resistance-associated kinome reprogramming, with AKT and p70S6K serving as central nodes in this network. Pharmacological inhibition of these kinases re-sensitized resistant cells to imatinib. Intriguingly, we identified LIN28A as a master regulator of this resistance kinase network: knockdown of LIN28A sensitized cells to imatinib whereas its overexpression conferred resistance. Our findings establish LIN28A-mediated kinome reprogramming as a novel mechanism of acquired TKI resistance and propose therapeutic targeting of the LIN28A-AKT/p70S6K/PKC axis to overcome treatment resistance in CML.

## EXPERIMENTAL PROCEDURES

### Cell Culture

K562 cells (ATCC, CRL-3344) were maintained in IMDM supplemented with GlutaMAX™ (Thermo Fisher Scientific, 31980030) and 10% fetal bovine serum (Wisent, 098-150) at 37°C in a humidified 5% CO_2_ atomosphere. To generate the imatinib-resistant subline (ImR), parental K562 cells were cultured in progressively increasing concentrations (0-5μM) of imatinib (Selleckchem, S2475) over a 6-month period. The established ImR line was maintained in medium containing 1μM imatinib. Prior to experiments, ImR cells were washed with PBS and cultured in drug-free medium for 72 hours to eliminate residual imatinib effects. For pharmacological studies, cells were treated with inhibitors to various kinases, including BCR-ABL: imatinib (S2475), dasatinib (S1021), or ponatinib (S1490); AKT: AKTi-1/2 (S7776); mTOR: rapamycin (Cayman, 13346); PKC/FLT3: midostaurin (S8064); SGK: GSK650394 (S7209); PDK1: BX-912 (Cayman, 14708; CaMKII: KN-93 (Cayman, 21472); or AXL: bemcentinib (MedChemExpress, HY-15150).

### Mass Spectrometry (MS) Analysis

Sample preparation for proteome and phosphoproteome analysis was performed as described previously^20^ with modifications. Protein samples (150 µg) was subjected to reduction with 5 mM Tris(2-carboxyethyl)phosphine (TCEP, BioShop) and alkylation with 15 mM iodoacetamide (IAA, Sigma-Aldrich). Proteins were concentrated using the single-pot, solid-phase-enhanced sample-preparation (SP3) method, followed by sequential digestion with LysC (1.5 mAU, Wako) for 2 h at room temperature and trypsin (1:50, wt/wt, Promega) for 16 h at 37°C. Peptide concentration was measured using the BCA assay (Thermo Fisher Scientific). For TMT-11plex labeling, 100 µg peptides per sample was resuspended in water (5 mg/mL) and labelled with 133 µg tandem mass tag (TMT) reagent (Thermo Fisher Scientific). To boost pTyr signal, the polled TMT mixture contained 20x spike-in of pervanadate-treated ImR sample (TMT 126 channel). After reserving 5% for proteome analysis, the TMT-labelled peptide mixture was incubated with SH2 superbinder agarose beads (Precision Proteomics, 001) for 1 h at 4°C to enrich pTyr peptides^21^. For pSer/pThr peptide enrichment, Ti^+4^-IMAC beads were applied to flow-through from SH2-superbinder enrichment. All samples (proteome, pY, and pS/pT) were fractionated using Pierce™ High pH Reversed-Phase Kit (Thermo Fisher Scientific) prior to LC-MS/MS analysis.

Peptides were analyzed using an UltiMate™ 3000 RSLCnano (Thermo Fisher Scientific) coupled to a Q-Exactive Plus mass spectrometer (Thermo Fisher Scientific) via an EASY-Spray ion source. The LC-MS system was operated in data-dependent acquisition mode. Peptides (in 0.1% TFA) were first loaded onto a PepMap Neo Trap Cartridge (5 mm 300 µm x 5 mm, Thermo Fisher Scientific 174500) and then separated on Double nanoViper PepMap Neo column (75 µm X 500 mm, 2 µm C18, DNV75500PN). The mobile phases comprise Buffer A: 0.5% acetic acid in water, and Buffer B: 0.5% acetic acid in 80% acetonitrile/water. For proteome and pS/pT peptide analysis, we used 300 nL/min flow rate with 4-hour and 3-hour gradients (from 100% Buffer A to 100% Buffer B), respectively. For pTyr peptide fractions, a 90-minute gradient was applied at 100 nL/min.

### MS Data Processing and Bioinformatic Analysis

Raw MS files were converted using MSConvertGUI^22^ and analyzed with Fragpipe (v20.0)^23^ using default settings for the TMT10-bridge workflow (proteome) and TMT10-phospho-bridge workflow (pS/pT and pY), modifying only the TMT interrogator setting from TMT10 to TMT11. The median-centered output was processed in R (version 4.2.2) for filtering, imputation, clustering, and statistical testing, with differential expression analysis performed using DEqMS^24^ for proteome (volcano plots) and LIMMA for phosphoproteomic data. Bioinformatic analysis included GSEA^25^ for proteome data and KSEA^26^ with phosphositePlus^27^ (downloaded 22/08/23) for phosphoproteome analysis. Protein-protein interaction networks were generated using STRING (confidence cutoff=0.7) via stringApp in Cytoscape (v3.8.0)^28^ used the stringApp and visualized with OmicsVisualizer^29^.

### Cell Proliferation and Viability Assays

Cell proliferation and viability were assessed using three complementary approaches. For WST-8 assays (Cayman), cells were plated at 100,000 cells/mL (200 µL/well in 96-well plates) and treated with inhibitor dose gradients; after 72 hours, 10 µL WST-8 reagent was added followed by 30 min incubation at 37°C before measuring absorbance (450 nm, background-corrected at 650 nm). Apoptosis was evaluated via Annexin V staining: cells (200,000 cells/mL in 6-well plates) were treated ± imatinib for 72 hours, washed with PBS, stained with APC-Annexin V (BioLegend) and Sytox AADvanced (Thermo Fisher) per manufacturers’ protocols, then analyzed on an LSRII flow cytometer (BD Biosciences) with FlowJo v10 software. For longitudinal proliferation tracking, cells (20,000 cells/mL) were stained with 32 nM Hoechst 33342 (30 min, 37°C), washed, resuspended in phenol red-free IMDM containing 3 µM propidium iodide, and imaged every 24 hours for 96 hours using a Cytation1 system (BioTek), with live cells (Hoechst+/PI-) quantified to generate growth curves analyzed in Prism v9.5.1 for doubling time calculations.

### Lentiviral Infection

Lentiviral particles were generated using third-generation self-inactivating vectors produced by transfecting HEK 293T cells with X-tremeGENE™ HP DNA Transfection Reagent (Roche) in serum-free Opti-MEM media^30^, using a plasmid mixture containing 0.5µg pMD2.G, 0.5µg pRSV-Rev, 1µg pMDLg/pRRE, and 0.5µg of the target plasmid. For knockdown experiments, we employed MISSION^®^ Lentiviral shRNA (Sigma) targeting AXL (TRCN0000001040), IGF2BP2 (TRCN0000255462), YBX1 (TRCN0000315309), and three-independent LIN28A constructs (TRCN0000021802, TRCN0000417094, TRCN0000416070). We used the Ef1a-LIN28-Ires-Puro plasmid (Addgene # 18921) to overexpress LIN28A. Sixteen hours post-transfection, the serum-free medium was replaced with DMEM containing 10% heat-inactivated FBS, followed by 48-hour incubation at 37°C. Viral supernatants were filtered through 0.45 µm PES syringe filters and used to transduce K562 or ImR cells in the presence of 10 µg/mL polybrene. After 72-hour incubation, transduced cells were either selected with 10 µg/mL puromycin (knockdowns) or sorted for GFP expression using FACSAria III (overexpression).

### RT-qPCR analysis

RNA extraction, reverse transcription, and quantitative PCR were performed as previously described^31^. Total RNA (including miRNA) was extracted from cell pellets using the miRNeasy Tissue/Cells Advanced Mini Kit (Qiagen, 217604) and quantified by absorption at 260 nm on a Nanodrop 2000 spectrophotometer. For cDNA synthesis, 2 µg of total RNA was reverse transcribed using the High-Capacity cDNA Reverse Transcription Kit (Applied Biosystems, 4368814) with miRNA-specific primers for miRNA targets and random primers for mRNA. Quantitative PCR (qPCR) was performed on a CFX Connect (Bio-rad) or Quantstudio 3 (Applied Biosystems) system with PowerTrack SYBR Green (Applied Biosystems, A46110) and target-specific primers. Relative quantification was calculated using the ΔΔCT method^32^.

### Western Blotting

Cells were lysed in NP-40 buffer (1% NP-40, 1% SDS, 20 mM Tris pH 7.6, 150 mM NaCl, 100 mM NaF, 1 mM PSMF) containing phosphatase inhibitors, sonicated (5 seconds on/off pulses, 40 sec total, 35% amplitude), and centrifuged (20,000 x g, 10 min). Protein concentration was determined by BCA assay. Samples (20 µg) was denatured in Laemmli buffer (BioRad) at 95 °C for 5 minutes, separated by SDS-PAGE, and transferred to PVDF-PSQ membranes (Millipore) overnight at 30 V (4 °C). Membranes were stained with FCF green, destained (6.7% acetic acid, 30% methanol), and total protein was imaged (ChemiDoc™ MP, IR680 channel). After blocking (TBST + 2% fish gelatin, 45 min), membranes were incubated overnight at 4°C with primary antibodies against: LIN28A (Bio-Techne, AF3757), GAPDH (CST, #97166), PTEN (CST #13866), RICTOR (Bethyl Laboratories, A300-459A) GSK3-α/β (Santa Cruz, sc-7291), pGSK-α/β (CST, #9331), pan-AKT (CST, #2920); pAKT-T308 (CST, #9275), pAKT-S473 (CST, #4060), mTOR (CST, #2972), mTOR-S2448 (BioLegend, 610302) ERK1/2 (Santa Cruz, sc-514302), pEKR1/2-Y204 (Thermo Fisher, MA5-15174), PDK1 (CST, #3062), pPDK1-S241 (CST, #3438). Following five 5-min TBST washes, blots were incubated (45 min) with fluorescent secondary antibodies: goat anti-mouse StarBright700 (Bio-Rad), donkey anti-goat AlexaFluor680 (Thermo), or donkey anti-rabbit AlexaFluor800 (Thermo). Protein bands were visualized (ChemiDoc MP) and quantified (ImageLab software) with normalization to total protein.

### Colony formation unit (CFU) assay

For CFU assays, 1,000 cells were suspended in IMDM medium containing 10% FBS and 0.9% methylcellulose (R&D Systems, HSC001), with or without 0.5 µM imatinib. The cell suspension (1 mL/well) was dispensed into 24-well plates using a 16-gauge needle (triplicate wells per condition). Plates were incubated at 37°C with 5% CO2 for 21 days, after which colonies were enumerated by microscopic examination.

### Cell Cycle Analysis

Following 24-hour treatments, cells were collected, washed with PBS, and viability-stained using the LIVE/DEAD™ Fixable Far Red Dead Cell Stain Kit. Fixed overnight in 70% ethanol (-20°C), cells were subsequently washed and stained in PBS containing 100 µg/mL RNase A and 50 µg/mL propidium iodide (37°C, 30 min). Cell cycle profiles were acquired on a BD LSR II flow cytometer and analyzed using FlowJo software (v10.0).

### Statistical Analyses

All statistical analyses were performed using either GraphPad Prism (v9.5.1) or R (v4.1.2) with the ggplot2 package. Results are presented as mean ± SEM from at least three independent biological replicates. Statistical significance was determined using either Student’s t-tests or one-way ANOVA, as appropriate for each experimental design. Significance levels are denoted as follows: *p < 0.05, **p < 0.01, ***p < 0.001, ****p < 0.0001, with ns indicating non-significant results (p ≥ 0.05).

## RESULTS

Imatinib resistance is associated with broad changes in the proteome and phosphoproteome To investigate the mechanism of imatinib resistance, we developed a model system by culturing K562 cells in gradually escalating concentrations of imatinib over six months until resistance to 5 μM imatinib was attained. The resulting imatinib-resistant cell line, dubbed ImR, had a half maximal inhibitory concentration (IC50) of 3.7 μM, compared to 161 nM for the parental K562 cells, corresponding to a 23-fold increase in imatinib resistance as measured by WST-8 assay (Fig. 1A). Intriguingly, ImR cells also demonstrated an 11-fold resistance to dasatinib, a second-generation BCR-ABL TKI, and a 12-fold resistance to ponatinib, a third-generation TKI (Fig. 1B,C). Subsequently, we conducted experiments to determine if imatinib resistance led to changes in apoptosis and/or cell cycle progression. Cell death was confirmed by assessing early and late apoptosis using annexin V and 7-AAD, respectively. While no significant change in cell death was observed in ImR cells when cultured with or without a maintenance concentration (1 μM) of imatinib, a significant decrease (from 88.2% to 11.0%) in live cells was detected in K562 in the presence of imatinib. Additionally, exposure of K562 cells to imatinib led to increases in both early (from 9.1% to 25.9%) and late (from 2.6% to 49.7%) apoptosis (Fig. S1A). Moreover, imatinib treatment of K562 led to a significant increase (from 24% to 38%) in cells in the G0/G1 phase, compared to ImR cells that exhibited no significant change in the cell-cycle with or without imatinib (Fig. S1B). Compared to K562, ImR cells demonstrated significantly greater colony-forming capacity in methylcellulose CFU assays. While K562 cells failed to form any colonies in the presence of imatinib, ImR cells not only maintained robust colony formation but also preserved their progenitor/stem cell population despite BCR-ABL inhibition (Fig. S1C). Therefore, compared to parental K562 cells, the imatinib-resistant derivative ImR cells exhibit enhanced proliferative capacity, increased colony-forming potential, and reduced susceptibility to apoptosis.

**Figure 1.**
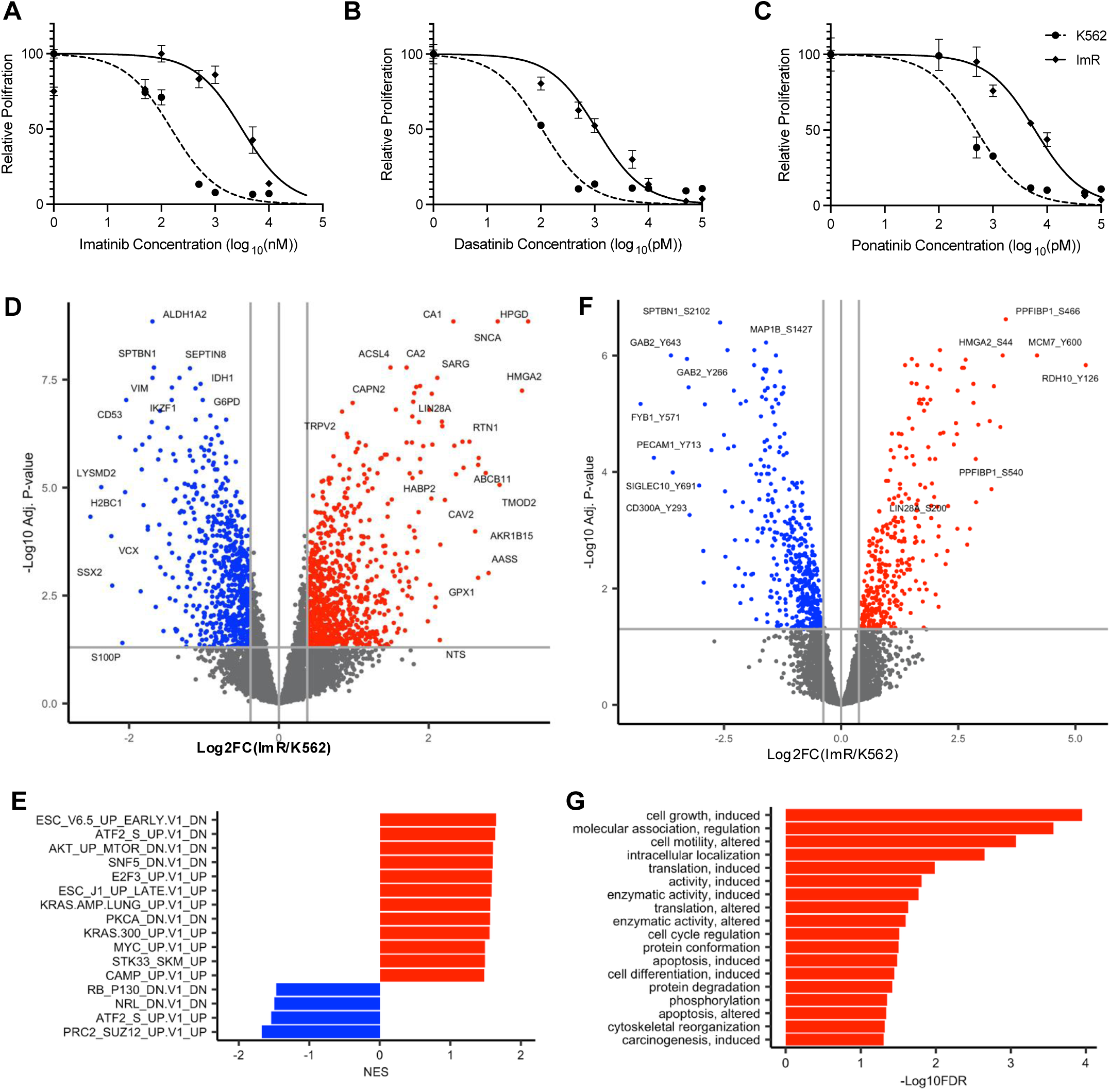
Proteome and phosphoproteome alterations associated with Imatinib resistance. (**A-C**) The ImR clone exhibited resistance to imatinib (**A**), dasatinib (**B**), and ponatinib (**C**), as demonstrated by relative proliferation of K562 and ImR cells following 72 hr inhibitor treatment (mean ± SEM, n=3 biological replicates). (**D**) Volcano plot illustrating differential protein expression between ImR and parental K562 cells. (**E**) GSEA revealed oncogenic signatures enriched in ImR (red) or K562 (blue) cells. (**F**) Volcano plot showing significantly increased (red) or decreased (blue) phosphosites in ImR compared to K562. (**G**) PhosphoSitePlus (PSP) regulatory site enrichment analysis of upregulated phosphosites in ImR cells. For all volcano plots, significance thresholds were set at a Benjamini-Hochberg (BH)-corrected p-value < 0.05 and minimum fold change of 1.3. All data were obtained from cells cultured in drug-free conditions.

To elucidate the molecular mechanism underlying imatinib resistance^33^, we performed quantitative mass spectrometry (MS) enabled by tandem mass tag (TMT) labelling to compare global proteomic and phosphoproteomic profiles between K562 and ImR cells (Fig. S1D). Our analysis identified 7,352 across all samples (Fig. S1D), with 1,550 showing significant differential expression (p<0.05) between cell lines (Fig. 1D). To account for ratio compression inherent in TMT-MS2 quantification^34^, we applied a 1.3-fold change (absolute value) for defining differentially expressed proteins (DEPs) and differentially phosphorylated proteins (DPPs). Using these criteria, we identified 879 upregulated and 671 downregulated proteins in ImR compared to K562 cells (Fig. 1D). Gene Set Enrichment Analysis (GSEA) of DEPs revealed significant enrichment of multiple oncogenic signatures and hallmarks in ImR cells (Fig. 1E; Fig. S2)^25^. Notably enriched pathways included those mediated by AKT-mTOR, KRAS, MYC, PKCA (PKCα), and ESC (embryonic stem cells) signaling. While the precise underlying mechanisms require further investigation, these pathways have been previously implicated in conferring TKI resistance in CML^35–37^.

To investigate whether imatinib resistance is associated with oncogenic signalling alterations, we compared phosphoproteomic profiles of ImR and K562 cells. Using Ti4+ immobilized metal affinity chromatography (Ti-IMAC) for pSer/pThr peptide enrichment combined with SH2 superbinder-based pTyr peptide enrichment, we identified 11,399 pSer/pThr sites and 1,103 pY sites^21,38,39^. After rigorous filtering across plexes, we quantified 6,511 pSer/pThr and 818 pY sites. Remarkably, 1,808 phosphosites were significantly altered between K562 and ImR cells (Fig. 1F). PhosphoSitePlus analysis demonstrated that ImR cells exhibited elevated phosphorylation at regulatory sites within critical pathways controlling cell proliferation, translation, cell cycle progression, motility, and apoptosis (Fig. 1G; Fig. S3)^27^. These quantitative proteomic and phosphoproteomic findings collectively demonstrate that imatinib resistance involves systematic reprogramming of both protein expression and phosphorylation-dependent signaling networks.

### Kinome Reprogramming Underlies Imatinib Resistance

Our mass spectrometry analysis detected 310 protein kinases, including 285 Ser/Thr kinases (STKs) and 25 Tyr kinases (TKs), with 49 showing significant elevation in either expression or phosphorylation levels in ImR cells (Fig. 2A, B; Fig. S4). Notably, the Tyr phosphoproteome profile clearly distinguished ImR from K562 cells, with 9 TKs demonstrating significantly increased expression and/or phosphorylation in ImR cells regardless of imatinib treatment (Fig. S4A, B). Among these altered TKs, AXL, EPHB6, and BTK showed concurrent increases in both protein expression and phosphorylation in ImR cells, strongly implicating these kinases in imatinib resistance (Fig. 2C). In contrast, phosphorylation of Tyr393 in ABL1’s activation loop and Y694 in STAT5A - a key downstream effector of ABL1 signaling - were significantly reduced in ImR cells, confirming effective inhibition of the BCR-ABL-STAT5 axis by imatinib (Fig. 2C). The kinome alterations extended beyond TKs, with 40 STKs showing significant upregulation in expression and/or phosphorylation. This included ARAF, a downstream effector of RAS signaling (Fig. 2E). These comprehensive kinome changes demonstrate that global reprogramming of both STK and TK networks represents a fundamental mechanism of imatinib resistance.

**Figure 2.**
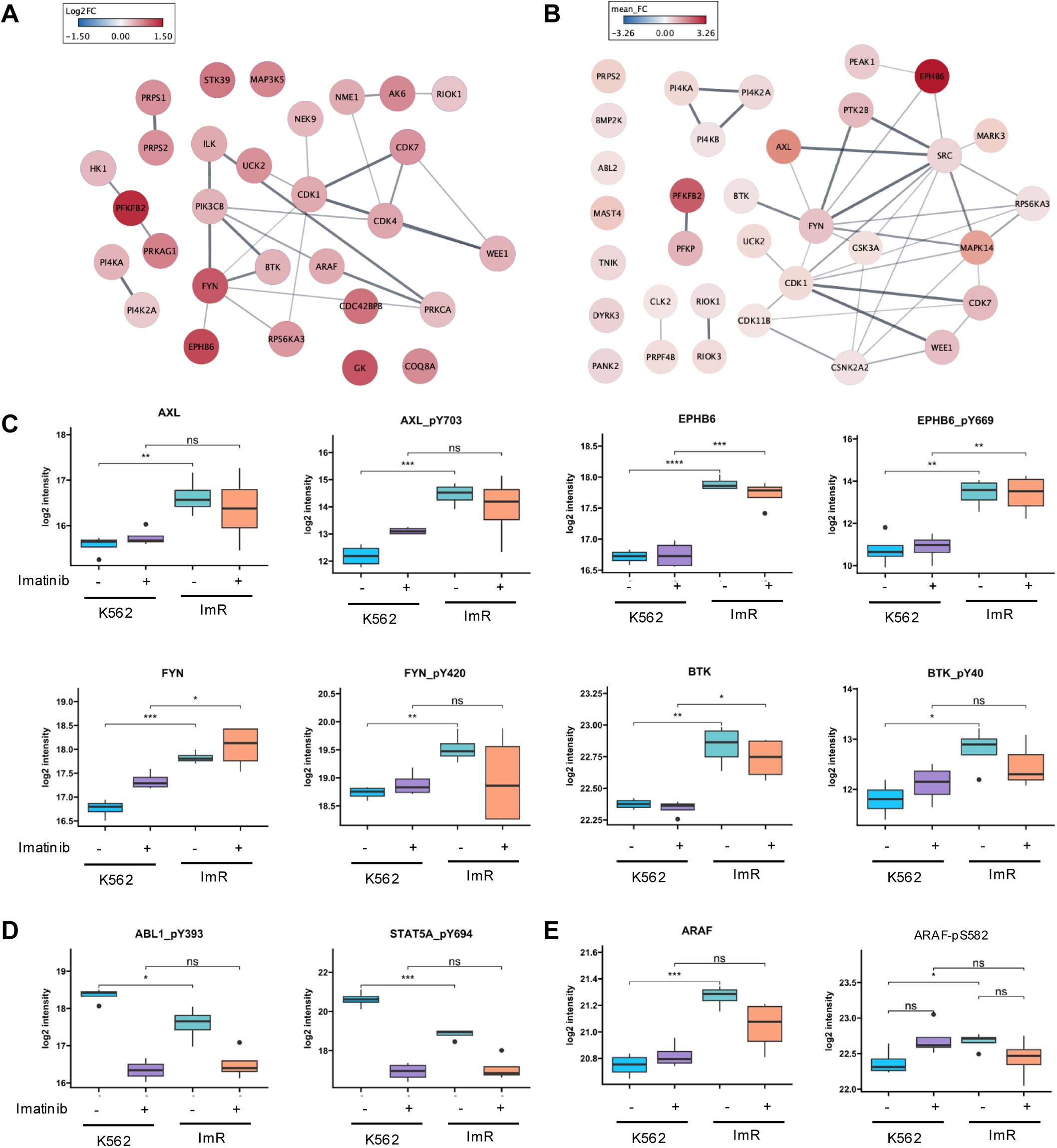
Kinome reprogramming in Imatinib-resistant cells. (**A**) Protein-protein interaction network of kinases significantly over-expressed in ImR compared to K562 at the proteome level. (**B**) Interaction network of kinases showing significantly increased phosphorylation in ImR versus K562, with collapsed fold-changes of detected phosphosites. Networks were created using StringApp in Cytoscape, with node color intensity representing fold-change and edge thickness indicating interaction confidence score. (**C**) Box plots showing representative TKs with a significant increase in both abundance and activity-regulatory site phosphorylation in ImR relative to K562. (**D**) Box plots demonstrating significant suppression of the ABL1-STAT5A signaling axis by imatinib treatment. (**E**) Box plots showing activation of the BCR-ABL downstream effector A-Raf in ImR cells. Statistical significance: * p<0.05; ** p<0.01, *** p<0.001 (Student t-test).

### A signalling network nucleated by AKT and p70S6K mediates imatinib resistance

While the activation status of most tyrosine kinases (TKs) can be gauged by phosphorylation levels of key regulatory sites, including activation loop or inducible Tyr sites, the regulation of many Ser/Thr kinases (STKs) involves different mechanisms. To gauge STK activity, we employed KSEA and Kinase Library (KL) to predict kinase activity based on phosphorylation levels of the corresponding substrates determined by MS^26,38^. Both programs predicted significant activation of the cell survival kinases AKT1 and AKT2 and the ribosomal protein kinases RPS6KA1/RSK1, RPS6KB1/p70S6K, RPS6KA3/RSK2 in ImR compared to K562 cells (Fig. 3A & B). This prediction was substantiated by the elevated phosphorylation of corresponding substrates in ImR (Fig. 3C-E). Other significantly activated STKs in ImR, identified through KSEA and/or KL, included SGK1 and SGK3, pivotal in stress response, PRKAA1/AMPK1 and PRKACA/PKACA, involved in energy-consumption, and PRKCD/PKCδ, a diacylglycerol-dependent kinase with diverse functions^36,39,40^. In contrast, the cytoplasmic tyrosine kinases (CTKs) HCK, FYN, SRC, LCK, LYN, BCR/ABL, and SYK were all significantly suppressed according to KSEA (Fig. 3A & B). These findings suggest that imatinib effectively inactivated CTKs while promoting the activation of numerous STKs.

**Figure 3.**
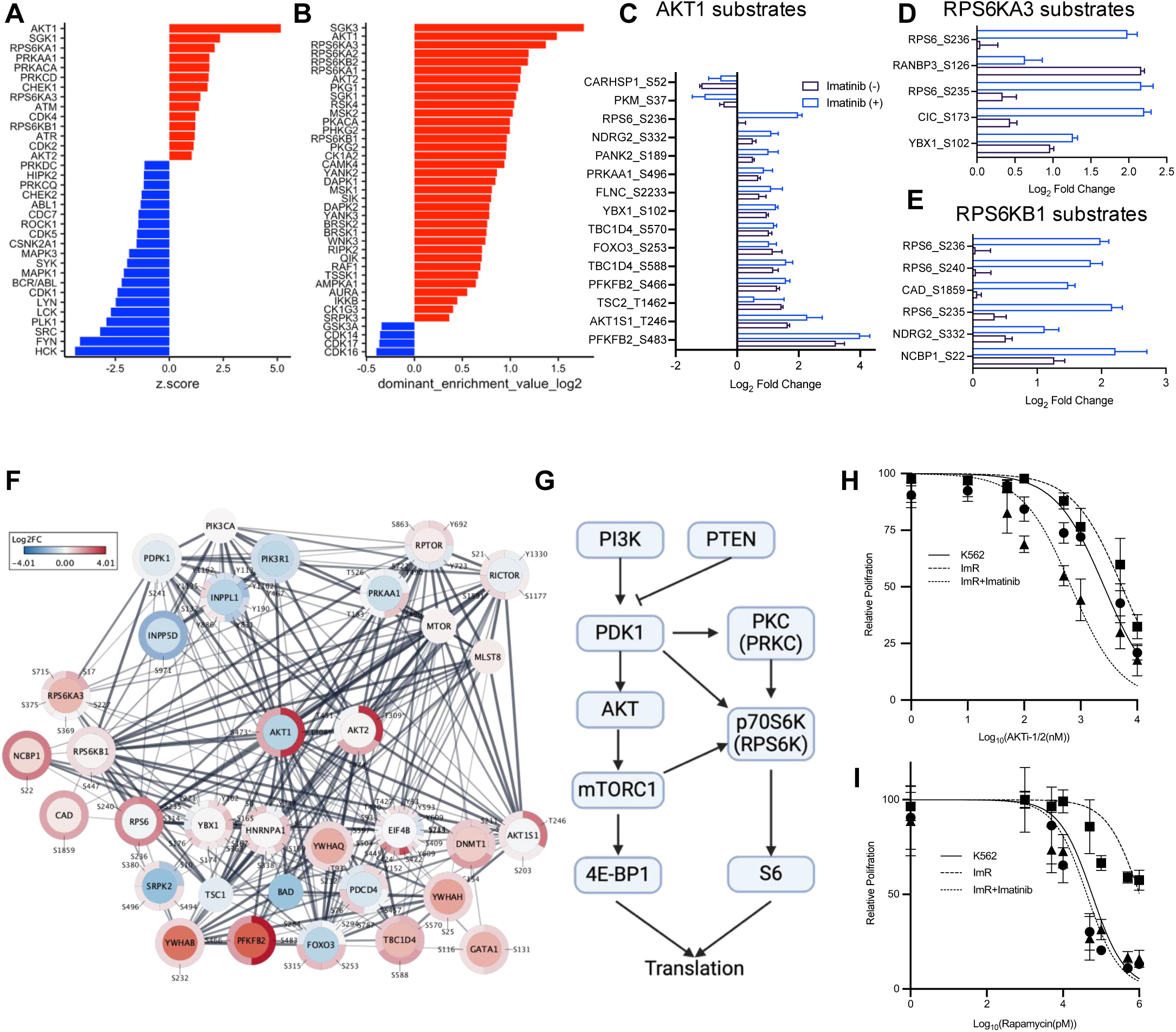
A signaling network nucleated by AKT and p70S6K underlies imatinib resistance. (**A, B**) Kinase activity predictions for (A) STKs/TKs by KSEA and (B) STKs by Kinase Library analysis. Activated kinases in ImR relative to K562 are shown in red, suppressed kinases in blue. (**C-E**) Substrates of AKT1, RPS6KA3, or RPS6KB1 demonstrating phosphoproteomic changes (log_2_ fold change >1 or <-1) between ImR and K562 cells. (F) Protein-protein interaction network centered on AKT1/AKT2 with upstream regulators and downstream effectors, constructed using Omics Visualizer and StringApp in Cytoscape. Inner and outer circles display protein expression and phosphorylation changes (ImR vs K562), respectively, with line thickness representing interaction confidence. Asterisks indicate phosphosites validated by Western blot. (G) Schematic of AKT-p70S6K (RPS6K) signaling pathways promoting protein translation. (H-I) Treatment with either (H) AKT1/2 inhibitor (ATKi-1/2) or (I) rapamycin, alone or combined with 100 nM imatinib, significantly restored imatinib sensitivity in ImR.

Our quantitative MS data revealed an extensive kinase-substrate interaction network within the PI3K-AKT-mTOR pathway, centered on AKT1, AKT2, and p70S6K/RPS6KB1, that likely drives imatinib resistance by enhancing survival and proliferation in ImR cells (Fig. 3F). Given the established role of AKT and p70S6K in protein translation (Fig. 3G), we investigated whether their pharmacological inhibition could restore imatinib sensitvity. Dual inhibition of AKT1/2 by AKTi-1/2 synergized with imatinib, reducing its IC50 by 7.9-fold in ImR cells (Fig. 3H). The combination showed maximal synergy at 1 μM (synergy score: 24.2, Fig. S5A). Strikingly, rapamycin (an mTORC1 inhibitor) enhanced imatinib sensitivity even more potently, lowering the IC50 by 24.4-fold and restoring it to levels comparable with K562 cells (Fig. 3I).

We further probed upstream regulators of AKT activation, including AXL, PDPK1 (PDK1, which phosphorylates AKT on T308), and CaMKII (which can phosphorylate AKT on S473). Bemcentinib (AXL inhibitor) modestly reduced imatinib IC50 (1.64-fold, Fig. S5B) while BX-912 (PDPK1/PDK1 inhibitor) showed no significant effect (Fig. S5C)^41,42^. KN-93 (CaMKII inhibitor), in contrast, decreased IC50 by 1.9-fold (Fig S5D)^43,44^. Notably, AKT pathway inhibitors affected both K562 and ImR cells similarly, consistent with AKT’s fundamental role in cell survival. These findings position AKT-mTOR signaling as a central node in imatinib resistance while highlighting mTORC1 inhibition as the most effective strategy to restore drug sensitivity.

### LIN28A plays an important role in mediating imatinib resistance

Given the established association between elevated LIN28A levels and cancer progression, we explored its potential role in leukemia drug resistance^16,19^. Analysis of available public clinical datasets revealed that high LIN28A expression correlates with poorer survival outcome in multiple cancers, including acute myeloid lymphoma (AML) (Fig. S6). Our quantitative MS data revealed significant overexpression of LIN28A in ImR cells compared to K562, along with increased phosphorylation at Ser200 (Fig. 4A). It has been shown in a previous study that phosphorylation of Ser200 by Erk1/2 promotes LIN28A’ let-7 independent activity^50^. Consistent with activation of let-7-independent mechanisms, we observed elevated expression or phosphorylation of multiple LIN28A-interacting proteins in ImR cells, including IGFBP2, YBX1, PRMT1, and PRMT5 (Fig. 4A; Fig. S7A, B). Paradoxically, we also found evidence for let-7-dependent pathway activation: DIS3L2 (a let-7-targetting exoribonuclease)^51^. was significantly upregulated in ImR cells growth without imatinib (Fig. 4B), accompanied by decreased levels of most let-7 miRNA family members (except let-7b) (Fig. 4C) and increased expression of let-7 target proteins (Fig. 4D). No significant changes were detected in LIN28B expression or phosphorylation (Fig. S7A). These findings demonstrate concurrent activation of both let-7-dependent and -independent LIN28A pathways in ImR cells, highlighting LIN28A’s central role in mediating imatinib resistance through multiple regulatory mechanisms.

**Figure 4.**
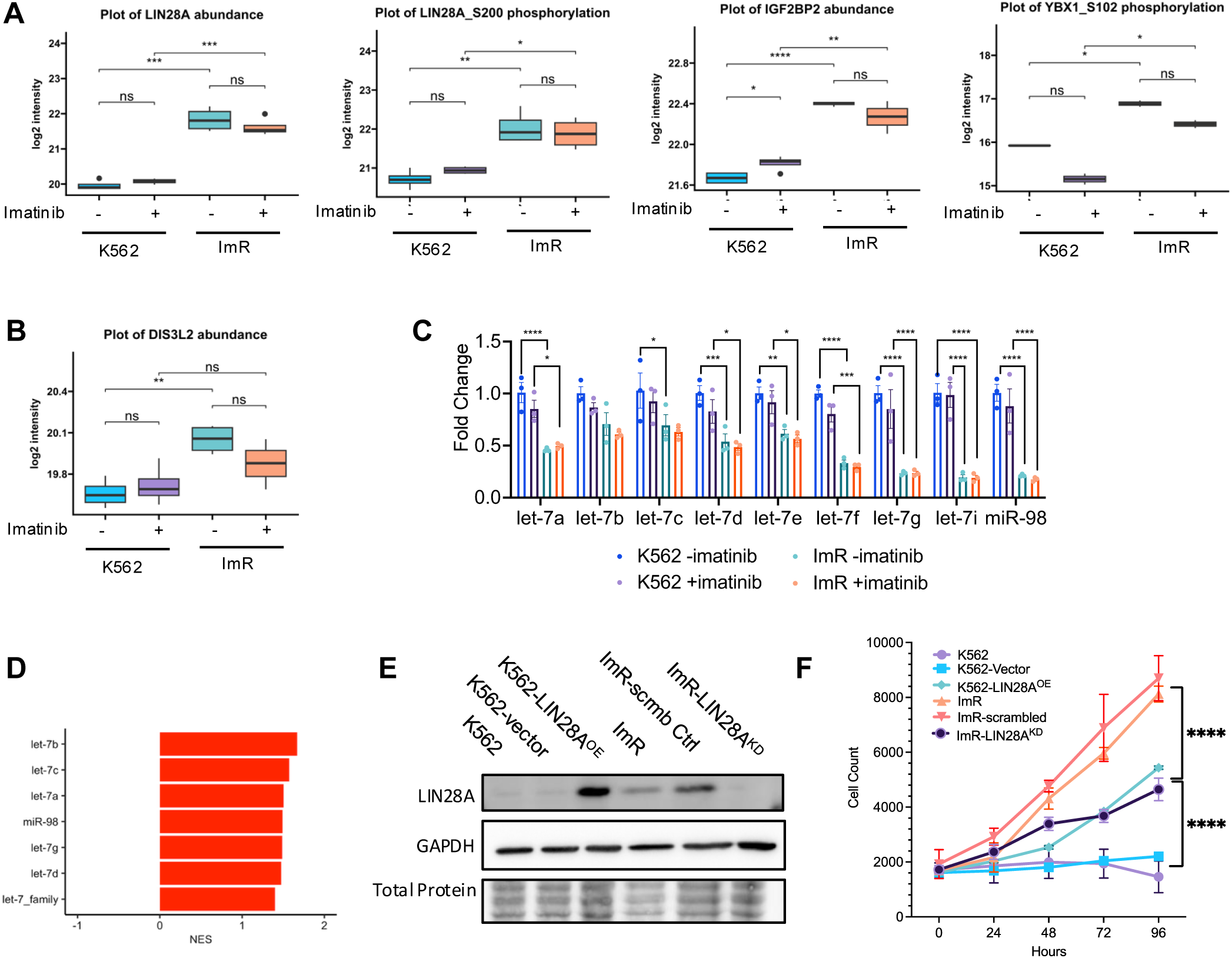
Essential role for LIN28A in imatinib resistance. (**A**) Proteomic and phosphoproteomic analyses reveal altered expression and phosphorylation of LIN28A interactors/regulators in the let-7-independent pathway. (**B**) Significant upregulation of DIS3L2, a negative regulator of let-7, in ImR cells. (**C**) RT-PCR analysis demonstrating reduced abundance of let-7 family miRNAs in ImR cells compared to K562, with or without 1 μM imatinib treatment (mean ± SEM; n=3 biological replicates). (D) Increased abundance of let-7 target proteins in ImR cells, as shown by GSEA normalized enrichment score. (**E**) Western blot validation of LIN28A overexpression in K562-LIN28A^OE^ cells and depletion in ImR-LIN28A^KD^ cells. (**F**) Growth curves showing cell proliferation in 500 nM imatinib over time (mean ± SEM; n=3-5 independent experiments). Statistical significance: *, p<0.05; **, p<0.01; ***, p<0.001; ***, p<0.001; ****, p<0.0001 (one-way ANOVA with Tukey’s post hoc test).

To investigate the temporal dynamics of resistance development, we analyzed earlier clones from the ImR selection process - including intermediate clones grown at 200 nM (Gen1) and 500 nM (Gen2) imatinib (Fig. S8A). Strikingly, we observed a stepwise increase in LIN28A expression levels that correlated with the progression of resistance, from parental K562 through Gen1, Gen2, and ultimately the ImR line (Fig. S8B-C). This increasing LIN28A abundance showed a strong positive correlation with rising imatinib IC50 values across the cell lines (Fig. S8D-E). Together with our previous findings, these results strongly suggest that LIN28A may play a functional role in driving imatinib resistance beyond mere association.

To establish a causal relationship between LIN28A and imatinib resistance, we performed LIN28A knockdown in ImR cells using shRNA (Fig. 4E, Fig. S9A, B). LIN28A depletion significantly impaired cell proliferation under imatinib treatment (1 μM), increasing doubling time compared to scramble controls (Fig. 4F; Fig. S9C). Strikingly, while control ImR cells maintained viability at 5 µM imatinib, LIN28A-depleted cells failed to survive, demonstrating LIN28A’s essential role in resistance (Fig. S9C). To assess whether LIN28A alone is sufficient to confer resistance, we transfected K562 cells with a LIN28A-expressing plasmid (Fig. 4E). The resulting LIN28A^OE^ cells demonstrated a significant increase in resistance to imatinib compared to the parental K562 cells (Fig. 4F). Nevertheless, the K562-LIN28A^OE^ cells did not attain the same level of imatinib-resistance as the ImR cells, suggesting that additional proteins, aside from LIN28A, may contribute to resistance. To explore this possibility, we used the corresponding shRNAs to knock down IGF2BP2 (Insulin-Like Growth Factor 2 mRNA Binding Protein 2) and YBX1 (Y-Box Binding Protein 1), both interactors of LIN28A, as well as AXL and CD44, implicated in therapeutic resistance in various cancers^16,50–53^. These four proteins all showed a significant increase in expression and/or phosphorylation in ImR cells (Fig. S8C; Fig. S10A). While the knockdown of IGF2BP2, AXL, or CD44 did not result in a significant change in imatinib resistance compared to scramble controls, knockdown of YBX1 led to a significant reduction in imatinib resistance (Fig. S10C, D).

To validate findings from our MS experiments, we performed western blotting analysis of LIN28A, ERK1/2 (an upstream kinase of LIN28A^48,54^), and AKT1 (which phosphorylates YBX1) (Fig. S11A). Consistent with MS data, LIN28A showed a 6.5-fold increase in ImR versus K562 cells, maintaining elevated expression during continuous imatinib treatment (Fig. S11A, B), indicating stable upregulation in resistant cells. While total ERK1/2 and AKT1 levels remained unchanged, their phosphorylation was significantly higher in ImR cells both with and without 24-hour imatinib exposure (Fig. S11C-E). Interestingly, acute imatinib treatment reduced ERK1/2 and AKT phosphorylation in ImR cells, revealing dynamic regulation of these kinases in imatinib resistance (Fig. S11). Taken together, these data indicate that the Lin28A-AKT/ERK1/2 axis contributes to imatinib resistance.

### LIN28A is a master regulator of cell growth and survival

The ImR cells displayed marked alterations in their phosphoproteome and proteome compared to K562. Given that LIN28A plays a pivotal role in regulating abundance of proteins, including kinases, through controlling mRNA and miRNA abundance, we hypothesized that variations in LIN28A levels would induce systematic shifts in protein expression and phosphorylation^59–61^. To investigate imatinib resistance at a systems level, we performed quantitative MS-based proteomic and phosphoproteomic analyses comparing K562 cells stably overexpressing LIN28A with empty vector controls. Indeed, LIN28A overexpression resulted in broad alterations in protein expression and phosphorylation, with 1,437 proteins exhibiting significant changes in abunance and 731 Ser/Thr/Tyr phosphorylation sites displaying altered phosphorylation (Fig. 5A, B). Notably, LIN28A upregulated multiple oncogenic pathways, including AKT-, KRAS-, and ESC-mediated signaling, as well as metabolic pathways, mirroring observations in ImR cells (Fig. 5A; Fig. S12). Phosphoproteomic analysis further revealed widespread changes in phosphorylation for proteins governing cell cycle, cell growth, apoptosis, and carcinogenesis (Fig. 5B; Fig. S13).

**Figure 5.**
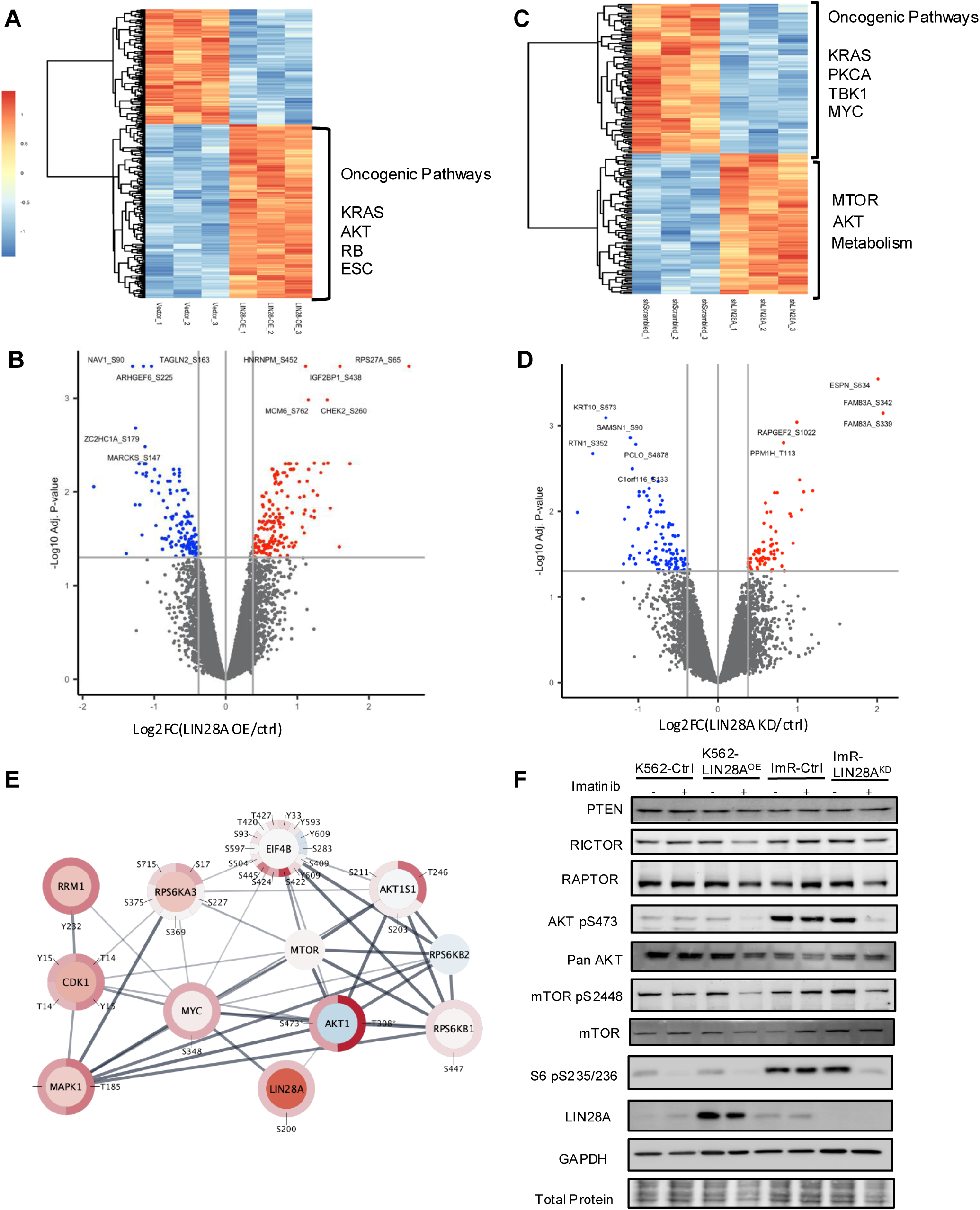
LIN28A regulates the protein translation machinery via AKT-RPS6K signaling. (**A**) Heatmap displaying differentially expressed proteins (DEPs) between K562-LIN28A^OE^ and empty vector control cells. (**B**) Volcano plot showing differentially phosphorylated proteins (DPPs) in K562-LIN28A^OE^ vs. control cells. (**C**) Heatmap of significant DEPs between ImR-LIN28^KD^ and control cells. (**D**) Volcano plot showing significant DPPs in ImR-LIN28^KD^ vs. control cells. (**E**) Interaction network centered on AKT, RSP6K, and LIN28A, with inner and outer circles representing DEPs and DPPs (Log2 fold change, ImR/ImR-LIN28^KD^) based on MS data. (**F**) Western blot analysis of AKT-mTOR pathway proteins in cells with LIN28A overexpression or knockdown, treated with 1 mM imatinib or vehicle for 24 hrs.

To validate these findings, we compared LIN28A-depleted ImR cells (i.e., LIN28A^KD^) with control ImR cells expressing scrambled shRNA to identify differentially expressed proteins (DEPs) and differentially phosphorylated proteins (DPPs). Depletion of LIN28A in ImR led to the downregulation of oncogenic pathways (e.g., KRAS, PKCA, and TKB) and the up-regulation of survival pathways (e.g., MTOR-AKT) along with metabolic reprogramming (Fig. 5C; Fig. S14). Similarly, DPPs were enriched for proteins regulating cell cycle, growth, protein translation, and oncogenesis (Fig. 5D; Fig. S15). Notably, through AKT1 and MYC, LIN28A is linked to a protein-protein interaction (PPI) network involving MAPK1 and ribosomal S6 kinases (Fig. 5E), which were differentially expressed and/or phosphorylated in ImR versus ImR-LIN28A^KD^ cells (Fig. 5E), suggesting that LIN28A may modulate cell proliferation and survival. Strikingly, this PPI network significantly overlaps with the one identified in ImR cells compared to K562 (Fig. 3F), reinforcing LIN28A’s central role in orchestrating adaptive responses to imatinib. Intriguingly, LIN28A depletion in ImR cells reduced AKT and mTOR phosphorylation without affecting their protein levels (Fig. 5F; Fig. S16), suggesting that LIN28A may regulate upstream kinases that activate these pathways^57,62^. Together, these findings highlight LIN28A as a key mediator of imatinib resistance by regulating adaptive changes in proliferation, survival, and metabolism.

### LIN28A orchestrates imatinib resistance by governing STK activity and signalling

To gain systematic insights into kinase regulation by LIN28A beyond ATK-mTOR, we performed kinase substrate enrichment analysis (KSEA) on phosphoproteomic data from K562-LIN28A^OE^ and ImR-LIN28A^KD^ cells. KSEA predicted elevated activity of multiple STKs in LIN28A^OE^ cells, including PKC family kinases (PRKCD/PKCδ, PRKCZ/PKCζ, PRKCA/PKCα), AKT2, stress-activated kinases (STK3, STK1, STK4, and SGK1, p70S6K (RPS6KA3), and CAMK2A. Notably, PRKCD/PKCδ and SGK1, which were hyperactive in ImR vs. K562 cells, were also upregulated in LIN28A^OE^ cells. Conversely, the LIN28A knockdown suppressed these kinases in LIN28A^KD^ cells (Fig.6A). Kinase library (KL) predictions further supported increased activity of PRKCD/PKCδ, PRKCA/PKCα, and AKT2 in K562-LIN28A^OE^ and ImR cells (Fig. S16). These findings were corroborated by corresponding changes in substrate phosphorylation patterns (Fig. S18-S19). Collectively, these results establish LIN28A as a master regulator of STKs, driving key signaling cascade that promote imatinib resistance in K562 cells.

**Figure 6.**
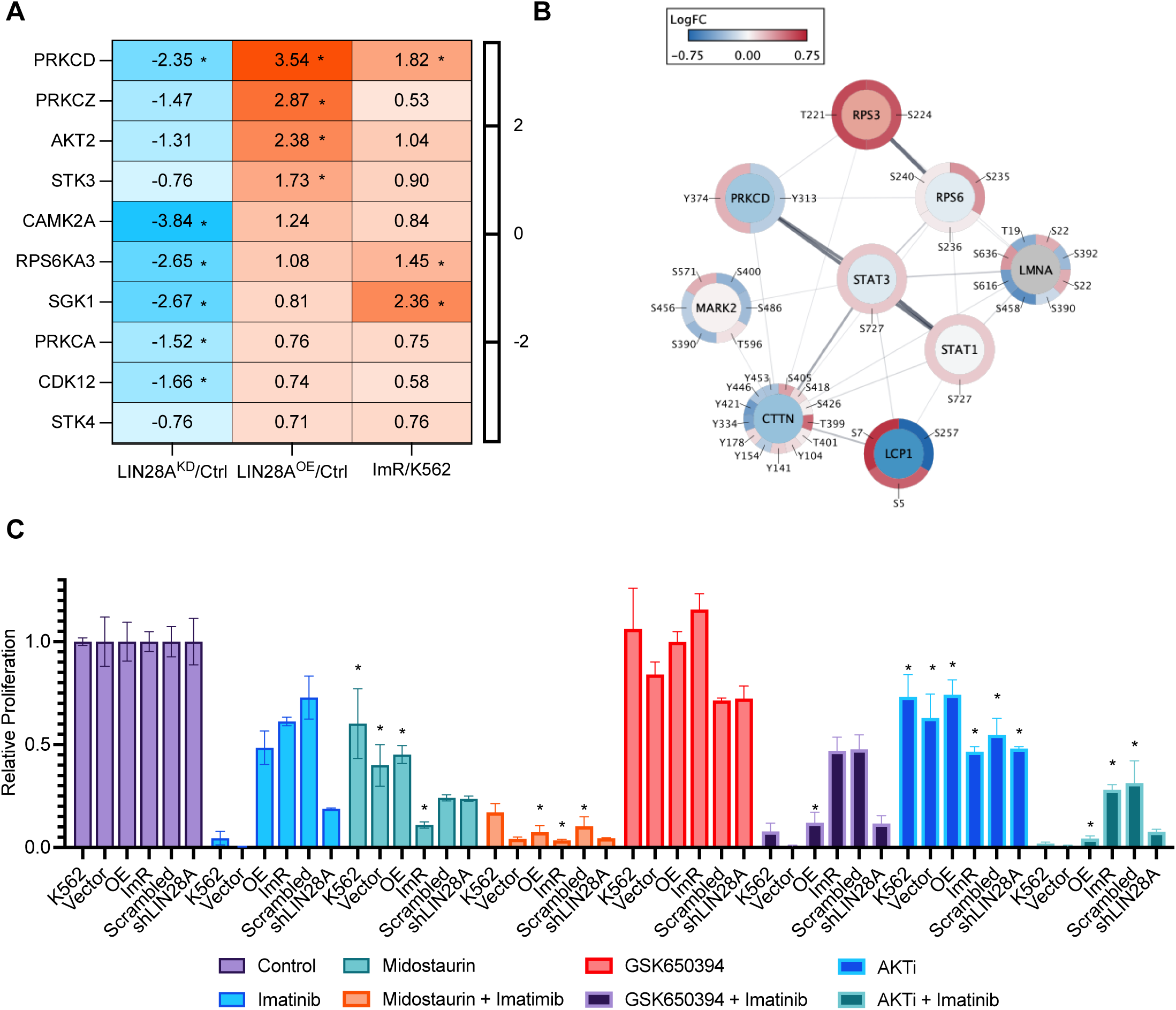
LIN28A-mediated reprogramming of an STK network promotes imatinib resistance. (**A**) Z-score of STKs predicted by KSEA to be activated (red) in ImR and K562-LIN28^OE^ cells and suppressed (blue) in ImR-LIN28A^KD^ cells. Asterisks (*) indicate significant prediction (FDR<0.05). (**B**) Signaling network centered on PRKCD (PKCd), showing differential protein expression (inner circle) and phosphorylation (outer circle) in ImR versus ImR-LIN28A^KD^ cells (Omics Visualizer and StringApp in Cytoscape). Color scale represents log2 fold changes from MS data. (**C**) Relative proliferation of the indicated cells treated with midostaurin (1mM, PKC inhibitor), GSK650394 (1mM, SGK1 inhibitor), or AKTi (1mM), with or without 500 nM imatinib. Cell viability was assessed by WST-8 assay after 72 hours. Data are normalized to vehicle (DMSO) or imatinib-only controls (mean ± SEM; *p < 0.05, one-way ANOVA with Tukey’s post hoc test).

In addition to AKT-mTOR, PRKCD/PKCδ emerges as a potential driver of imatinib resistance. Notably, PKCδ exhibited enhanced phosphorylation at Y313 and Y334 in ImR cells compared to K562 under imatinib treatment. Intriguingly, Src, a known upstream activator of PKCδ, was significantly upregulated in ImR cells (Fig. S20A), suggesting potential Src-PKCδ pathway activation^61–63^. Supporting this hypothesis, we observed increased phosphorylation of multiple PKCδ substrates, including ribosome proteins (RPS3 and RPS6), transcription factors (STAT3 and STAT1), and Src substrate (cortactin/CTTN) (Fig. 6B; Fig. S20). These findings collectively underscore PKCδ’s pivotal role in orchestrating imatinib resistance through an extensive signaling network.

### Targeting LIN28A-regulated kinases overcomes imatinib resistance

To evaluate the therapeutic potential of LIN28A-regulated kinases (PKC, SGK1, and AKT), we treated K562 cells (wild-type, LIN28A-overexpression, or empty vector control), ImR cells (wild-type, LIN28A-knockdown, or scrambled shRNA control), with specific inhibitors-either alone or in combination with imatinib. PKC inhibition (midostaurin) showed stronger anti-proliferative effects in ImR and ImR-LIN28A^KD^cells than in K562 or K562-LIN28A^OE^ cells. Combination with imatinib potentially suppressed proliferation of all cell types, with maximal efficacy in ImR cells (Fig. 6C). SGK1 inhibition (GSK650394), while exhibiting no standalone effect, significantly inhibited proliferation when co-administered with imatinib (Fig. 6C). Similarly, AKT inhibition (AKTi) enhanced imatinib’s efficacy, but was more toxic to K562 cells than to ImR cells (Fig. 6C). These results highlight PKC as a potent and selective target for overcoming imatinib resistance, with SGK1 and AKT as additional synergistic targets in the K562 model.

## DISCUSSION

While imatinib revolutionized CML treatment, resistance remains a significant challenge-particularly in blast stage patients^8^. Although second and third-generation TKIs (dasatinib, nilotinib, busotinib, and ponatinib) and the allosteric inhibitor sciminib target BCR-ABL kinase mutations^7,66,67^, resistance persists, underscoring the importance of BCR-ABL-independent mechanisms. Our quantitative proteomic and phosphoproteomic analyses of K562 versus imatinib-resistant (ImR) cells revealed that kinome reprogramming and associated phosphorylation signaling pathways drive resistance development. These findings highlight critical therapeutic targets beyond BCR-ABL mutations.

While imatinib effectively suppressed BCR-ABL activity, it paradoxically triggered activation of multiple tyrosine kinases (TKs), including AXL, EPHB6, and BTK (Fig. 2B). These TKs collectively activate MAPK and PI3K pathways^68^, creating redundant survival signals that render single TK-inhibition ineffective against resistance (Fig. 7)^69^. In contrast, targeting LIN28A-a MAPK/ERK substrate upregulated in ImR cells^50^- or its downstream Ser/Thr kinases (STKs), significantly impaired ImR proliferation. Notably, LIN28A depletion restored imatinib sensitivity in ImR cells whereas LIN28A overexpression induced resistance in K562 cells. Proteomic and phosphoproteomic analyses revealed that LIN28A orchestrates a kinome reprogramming centered on STKs (e.g., PKC, AKT), which critically regulate translation and survival. Pharmacological inhibition of these kinases—particularly PKC—potently suppressed ImR proliferation. These findings establish a model wherein LIN28A acts as a central hub, driving resistance through STK-mediated proliferation and survival pathways (Fig. 7).

**Figure 7.**
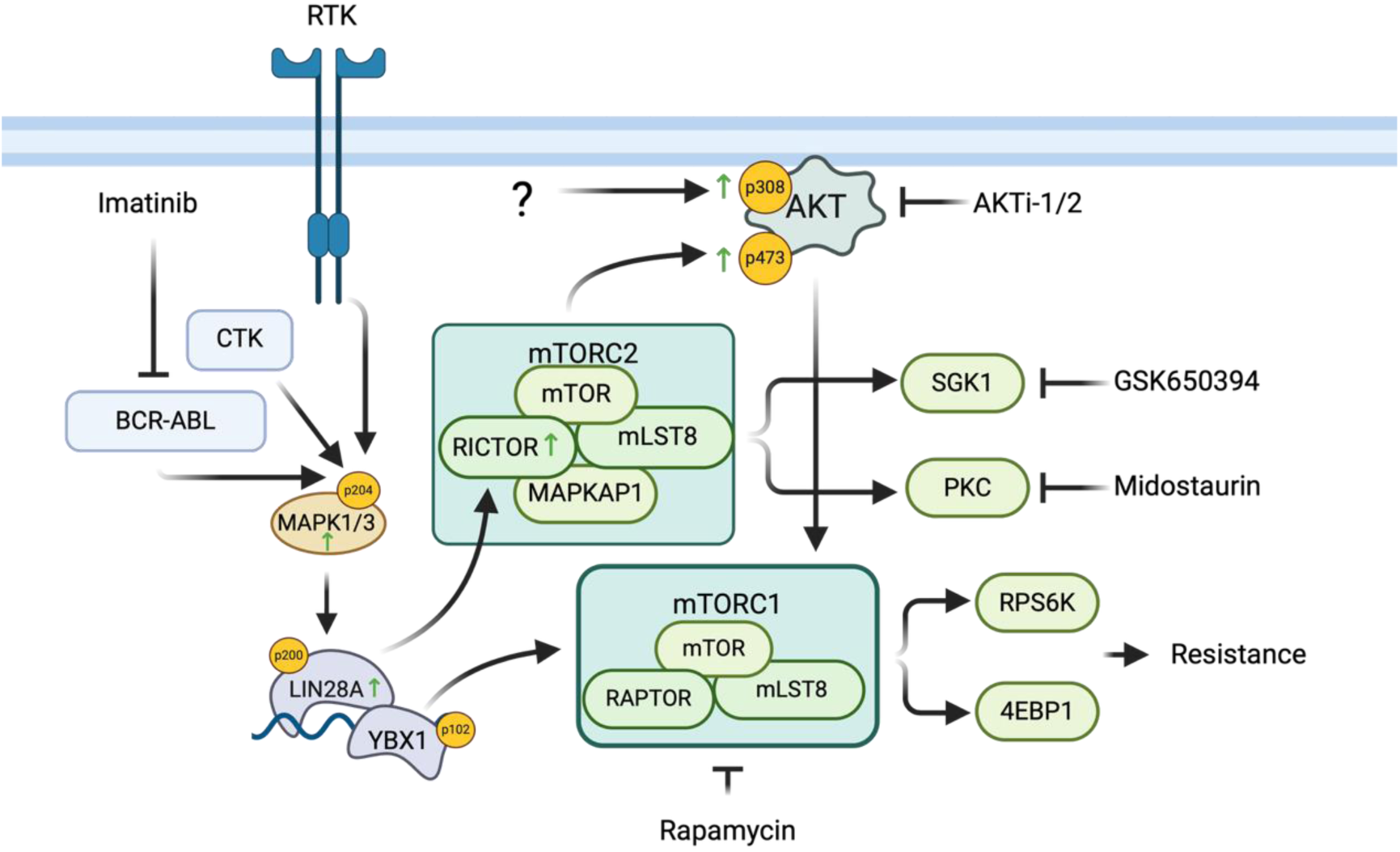
**Proposed model of LIN28A-regulated kinase signaling in imatinib resistance.**

The LIN28-let-7 axis play a well-established role in tumorigenesis across multiple cancers (e.g., breast, lung, ovarian, and pancreatic) and contributes to therapeutic resistance, including resistance to kinase inhibitors^16^. Normally a fetal gene, *LIN28A* is silenced in most adult tissues (except select muscle cells)^70^. Conditional LIN28A knockout shows no adverse effect in adult mice^71^. While LIN28A is expressed in approximately 9% of CML patients at the chronic phase, elevated expression is detected in 40% blast phase/blast crisis cases^19^. Intriguingly, its homolog LIN28B increases in 23% of chronic-phase patients progressing to blast crisis^19^. ADAR1 overexpression in blast crisis was found to promote LIN28B expression in a dasatinib-resistance model^72^. In a nilotinib-resistance model, FLT3-TAZ upregulation in blast crisis was shown to increase CD36 and LIN28A expression^73^. Though based on limited samples, these findings strongly implicate LIN28A/B in CML progression and highlight their potential as therapeutic targets.

LIN28A regulates cellular functions through both let-7-dependent and let-7-independent mechanisms. In the let-7-dependent pathway, LIN28A controls gene expression by promoting the degradation of let-7 family microRNAs, thereby modulating numerous oncogenes and tumor suppressors^74^. Our findings demonstrate that this LIN28A-let-7 axis is involved in imatinib resistance, as elevated expression of LIN28A in ImR cells correlates with reduced let-7 miRNA levels and concomitant upregulation of let-7 target genes. In parallel, LIN28A also operates through a let-7-independent mechanism where it associates with mRNA-stabilizing proteins including YBX1, PRMT1, PRMT5, and IGF2BP2 to enhance gene expression. Notably, these interacting partners show increased expression and/or phosphorylation in ImR cells, indicating activation of the let-7-independent pathway during resistance development (Fig. 7). While both pathways are clearly engaged in imatinib-resistant cells, their respective contributions to the observed STK reprogramming and consequent drug resistance remain to be fully elucidated. The relative importance of these two mechanisms in driving the kinase network remodeling that underlies therapeutic resistance represents a critical area for future investigation.

Our work identifies LIN28A as a promising therapeutic target for overcoming imatinib resistance. Several LIN28A inhibitors have recently been developed, including 1632, LI71, SB1301, C902, PH-43, TPEN, Ln7, Ln15, and Ln115^73–75^. Among these, compound 1632 has shown favorable safety profiles in multiple murine models^76,77^, warranting further investigation in ImR and other imatinib-resistant systems. These inhibitors could also serve as chemical scaffolds for developing LIN28A-targeted degraders using PROTAC technology^78^. An alternative strategy involves targeting LIN28A-regulated kinases, particularly PKC. Supporting this approach, midostaurin-an oral multiple kinase inhibitor of FLT3 and PKC, displayed potent anti-proliferative effects against ImR cells both as monotherapy or in combination with imatinib. This finding suggests the potential of repurposing midostautin, currently used for FLT3-ITD-positive acute myeloid leukemia, to treat blast stage or imatinib-resistant CML characterized by LIN28A and PKC hyperactivation^79^. Notably, midostaurin’s broad kinase inhibition profile (including CAMKK2, AURKA/B, RPS6KA, and PDK1^80^ may contribute to its efficacy against resistant cells by simultaneously blocking multiple LIN28A-regulated pathways beyond PKC alone. This polypharmacology likely explains its superior ability to resensitize ImR cells compared to more selective kinase inhibitors.

In summary, our study reveals LIN28A and its regulated kinase network as central players in imatinib resistance. We have not only delineated the mechanistic framework by which LIN28A-mediated kinome reprogramming drives therapeutic resistance but also identified PKC and AKT as key downstream effectors that could be targeted therapeutically. The observation that ImR cells show cross-resistance to second- and third-generation BCR-ABL inhibitors suggests this mechanism may represent a broader paradigm of TKI resistance in CML. Future investigations should validate these findings across diverse resistance models, including cells harboring clinically relevant mutations like T315I, and crucially, in primary samples from TKI-resistant CML patients. These studies will be essential for translating our discoveries into clinically actionable strategies to overcome treatment resistance.

## ACKNOWLEDGMENTS

We thank Kun Ping Lu, Xiao Zhen Zhou, and Patrick O’Donoghue for their valuable insights and critical discussions. This research was supported by grants from the Canadian Institute of Health Research (to SSCL), the Natural Sciences and Engineering Research Council of Canada (to SSCL and IUH); the Ontario Ministry of Research and Innovation (to IUH). OFJH received funding from Thermo Scientific Tandem Mass Tag Research Award which provided reagents for this project. OFJH held a Translational Breast Cancer Research Scholarship from the Canadian Cancer Society. MIF held a Queen Elizabeth II Graduate Scholarship in Science and Technology (QEII-GSST). SSCL holds the Canada Research Chair and Wolfe Medical Research Professorship in Molecular and Epigenetic Basis of Cancer.

## AUTHOR CONTRIBUTIONS

SSCL and OFJH conceived and designed the project. OFJH, MIF, QMQ, JHK, AW, KY, TW, XR, TM, and CV performed the research. OFJH, SSCL, and JUH analyzed the data. SSCL and OFJH wrote the manuscript with input from IUH and MIF.

## DATA AVAILABILITY

The proteomic data has been uploaded to the PRIDE database: http://www.ebi.ac.uk/pride

Project accession: PXD064276

