## Supplemental_Figures for "LIN28A-Dependent Kinome and Phosphoproteome Reprogramming Promotes Imatinib Resistance"

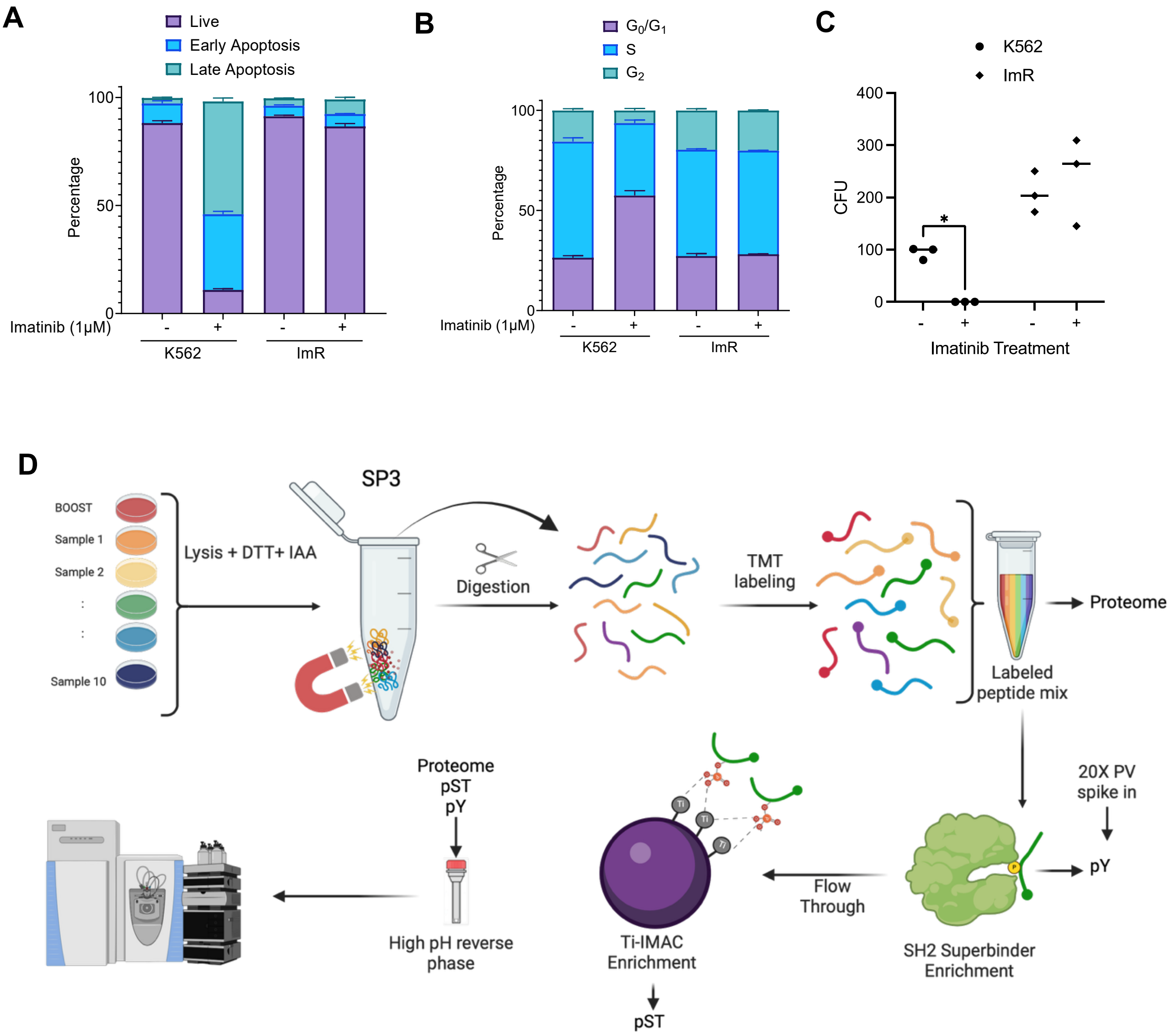


**Figure S1. Characterization of ImR, an imatinib-resistant clone of K562.** (**A**) Comparison of K562 and ImR cells for apoptosis using annexin V and 7-AAD staining. Cells were treated with 1 μM imatinib (+) or vehicle (-) for 72 hrs prior to staining. (**B**) Cell cycle analysis by flow cytometry. K562 and ImR cells were treated with and without imatinib for 24h, fixed and stained with propidium iodide, followed by flow cytometry analysis (mean ± SEM, n=3 biological replicates). (**C**) ImR cells formed more colonies than K562 based on methylcellulose CFU assay (14 days). *, $p<0.05$ (One-way ANOVA with Tukey post hoc test). (**D**) Workflow schematic for TMT mass spectrometry analysis.

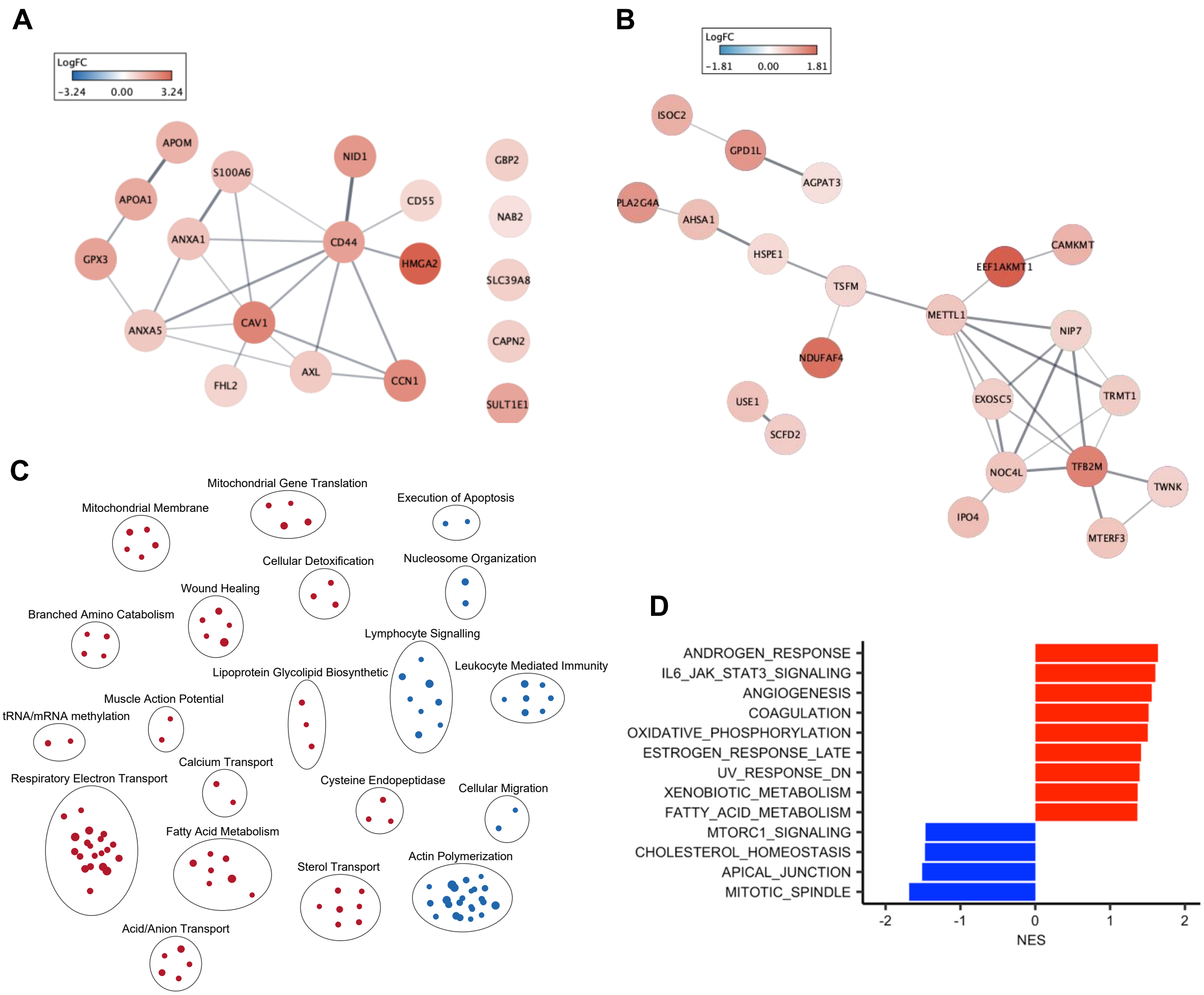


**Figure S2. Imatinib resistance involves rewiring of multiple signalling pathways.** (**A-B**) Interaction network of the leading edges of genes/proteins in the oncogenic signature “ESC_V6.5_UP_EARLY.V1_DN”+“ESC_J1_UP_LATE.V1_UP” (**A**), and ”MYC_UP.V1_UP” (**B**) (refer also to Fig. 1E). The circles are colored according to the log2FC(ImR/K562) ratio (scale bar). (**C**) EnrichmentMap of the enriched GO terms in ImR compared to K562. (**D**) Gene set enrichment analysis (GSEA) identified hallmark gene sets enriched in ImR (red) or K562 (blue). All analyses were based on the corresponding proteome data.

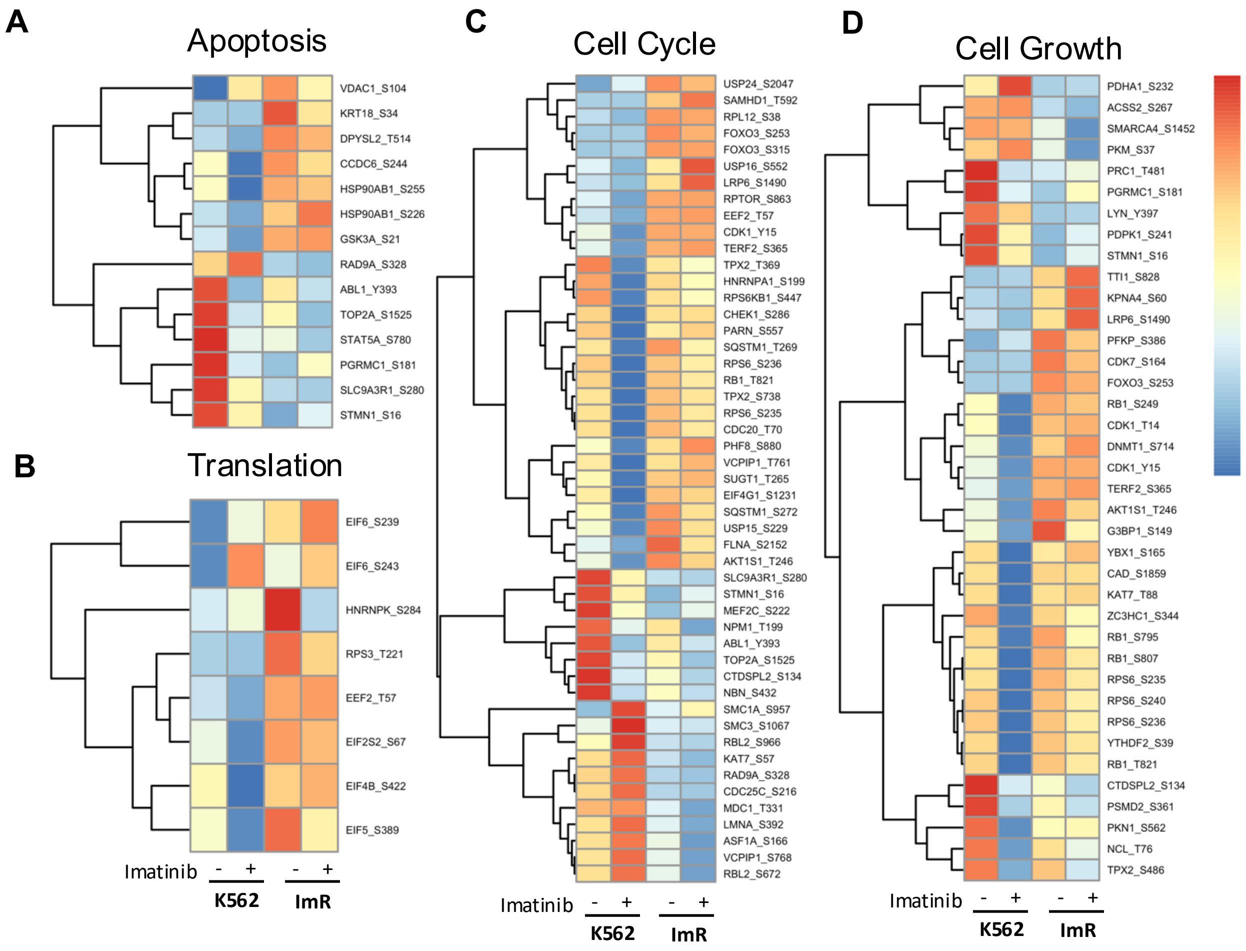


**Figure S3. Imatinib resistance alters phosphorylation signaling associated with apoptosis, translation, cell cycle, and cell growth.** In relation to Figure 1G, significantly altered phosphosites were clustered using Pheatmap based on the corresponding z scores. Heatmaps are shown for proteins involved in apoptosis (A), translation (B), cell cycle (C), and cell growth (D) identified from PSP regulatory sites, comparing K562 and ImR. +/- indicates cells treated with 1μM imatinib or vehicle for 24 hours.

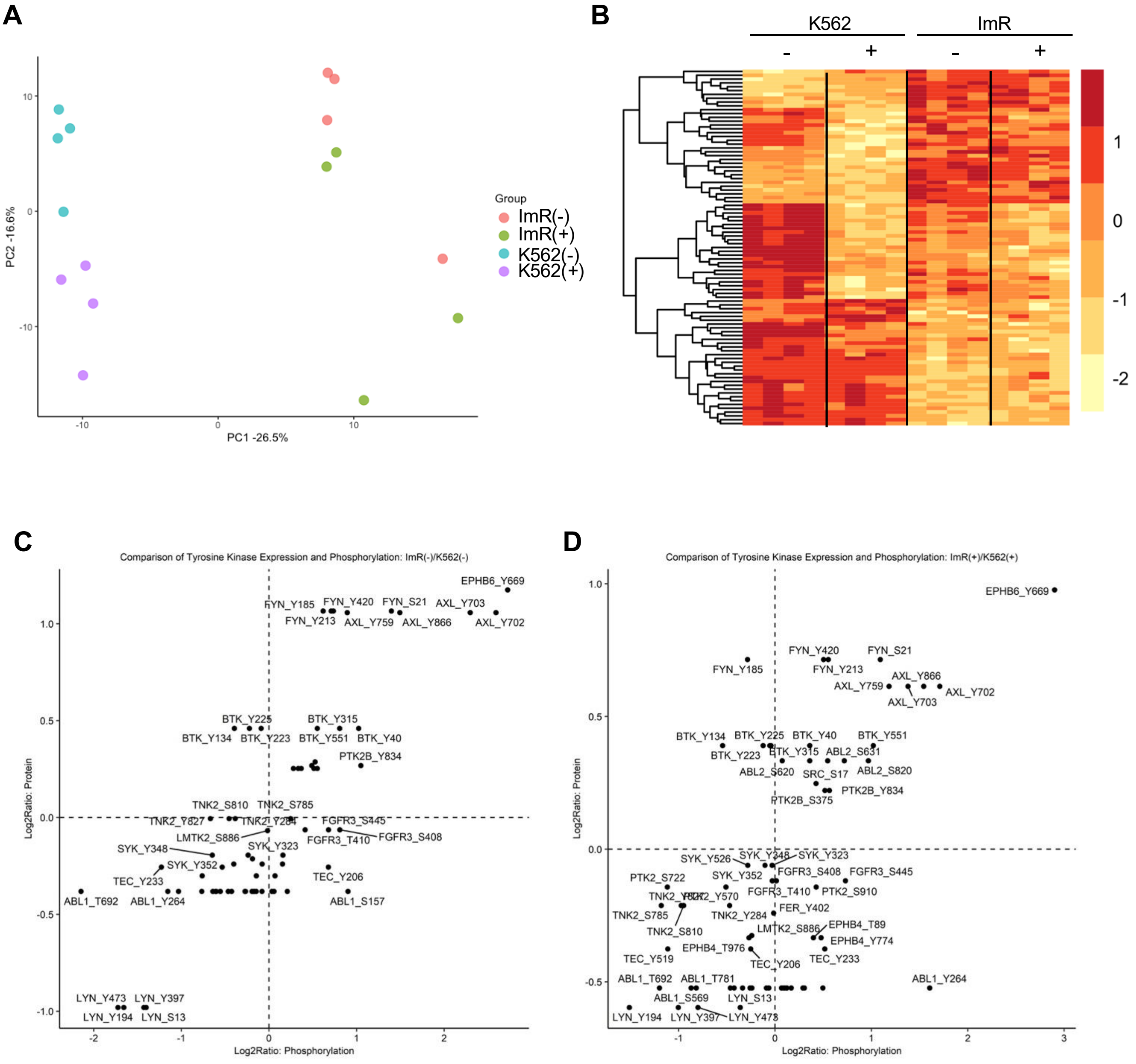


**Figure S4. Imatinib resistance is accompanied by alterations in tyrosine kinases and Tyr phosphorylation.** (**A**) PCA plot comparing tyrosine phosphorylation sites in K562 and ImR cells (+/- 1µM imatinib). (**B**) Heatmap showing differentiational phosphorylated tyrosine sites between ImR and K562. Cutoffs for heatmap were set to 0.05 BH-corrected p-value and 1.3 minimum fold change. (**C**-**D**) Tyrosine kinases differentially expressed and/or phosphorylated between ImR and K562 cells cultured in the absence (**C**) or presence (**D**) of 1 uM imatinib.

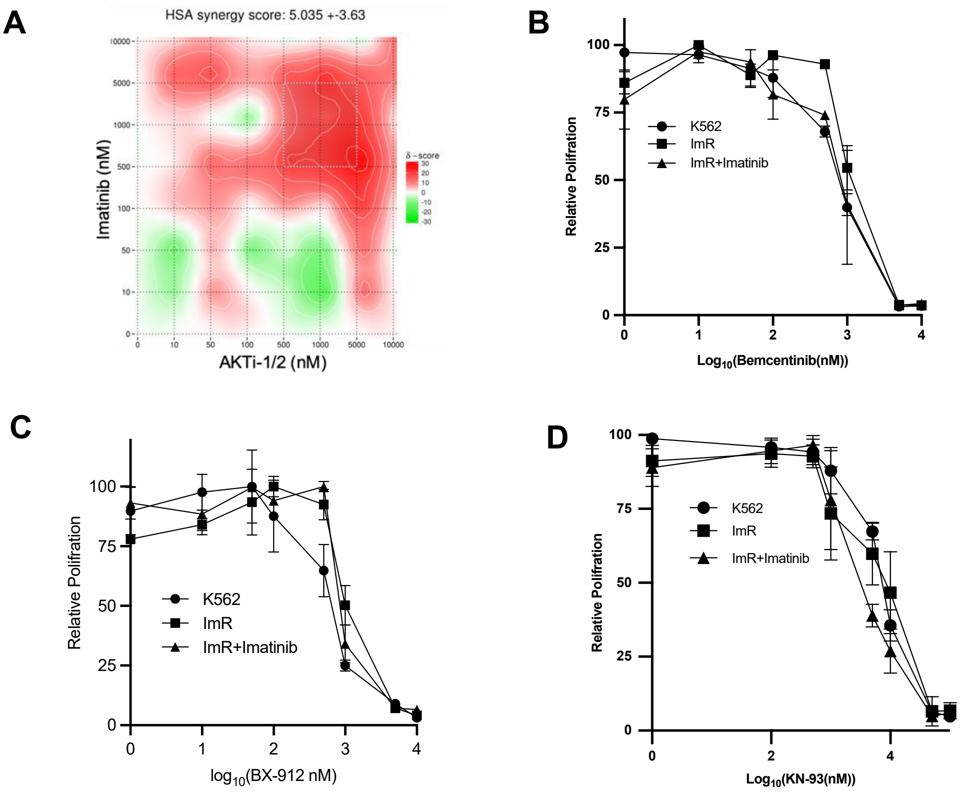


**Figure S5. Combination of imatinib and PI3K-AKT pathway inhibitors led to reduced proliferation in ImR cells. (A)** The combination of AKTi-1/2 and imatinib are synergistic, as assessed using SynergyFinder 2.0. (**B-D**) Pharmacological inhibition with the AXL inhibitor Bemcentinib (B), PDK1 inhibitor BX-912 (C) or CAMK inhibitor KN-93 (D) singularly or together with 100 nM of imatinib did not significantly restore sensitivity to imatinib.

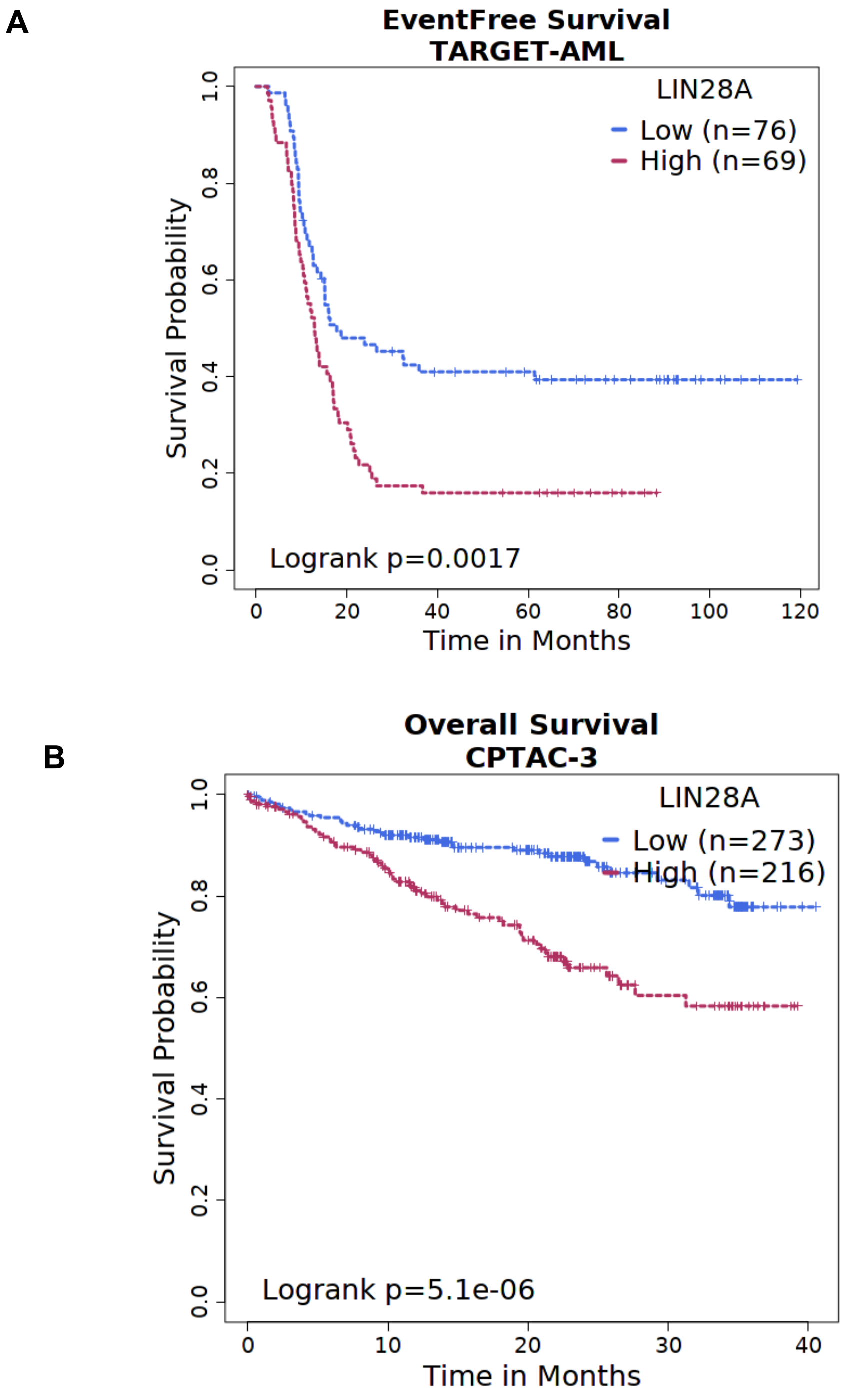


**Figure S6. Lin28A expression correlates with patient survival in multiple cancers.** Shown are Kaplan-Meier survival curves for (A) AML, based on Therapeutically Applicable Research to Generate Effective Treatments (TARGET) dataset, and (B) pan-cancer, based on datasets from Clinical Proteomics Tumor Analysis Consortium or CPTAC-3. LIN28A expression is inversely correlated with event-free or overall survival.

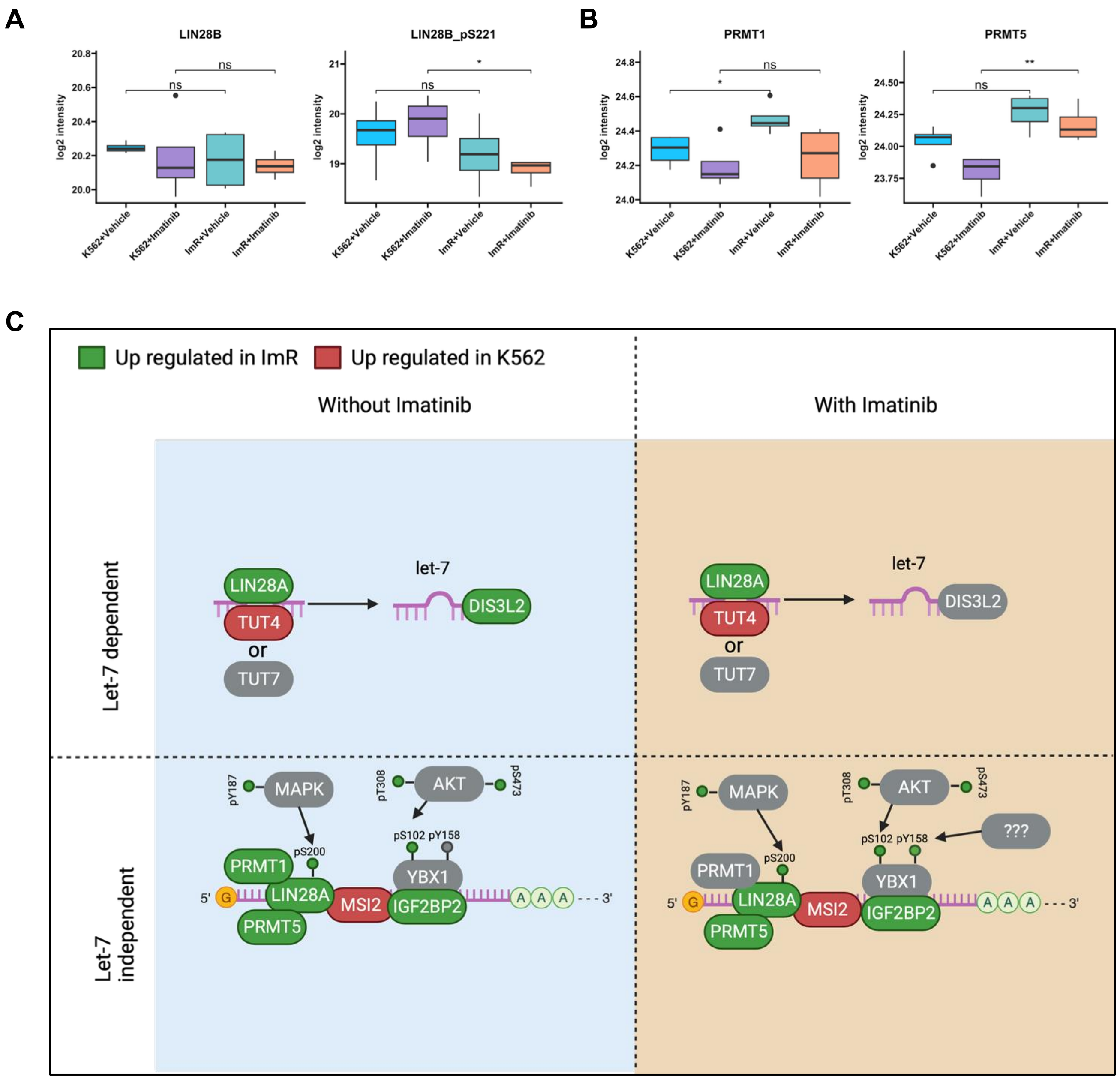


**Figure S7. Let-7-dependent and let-7-independent pathways of LIN28A in imatinib resistance.** (**A-B**) Box plots showing expression and phosphorylation of LIN28B (A) as well as PRMT1 and PRMT5 (B) in ImR compared to K562 cells. +/-, cells cultured with/without imatinib. *, p<0.05, ANOVA with Tukey post hoc test. (**C**) Schematic diagrams highlighting let-7-dependent (upper panel) and let7-independent (lower panel) LIN28A function. Proteins with increased expression or phosphorylation in ImR or K562 (-/+ imatinib) cells are denoted in green or red, respectively. Those without significant changes are indicated in grey.

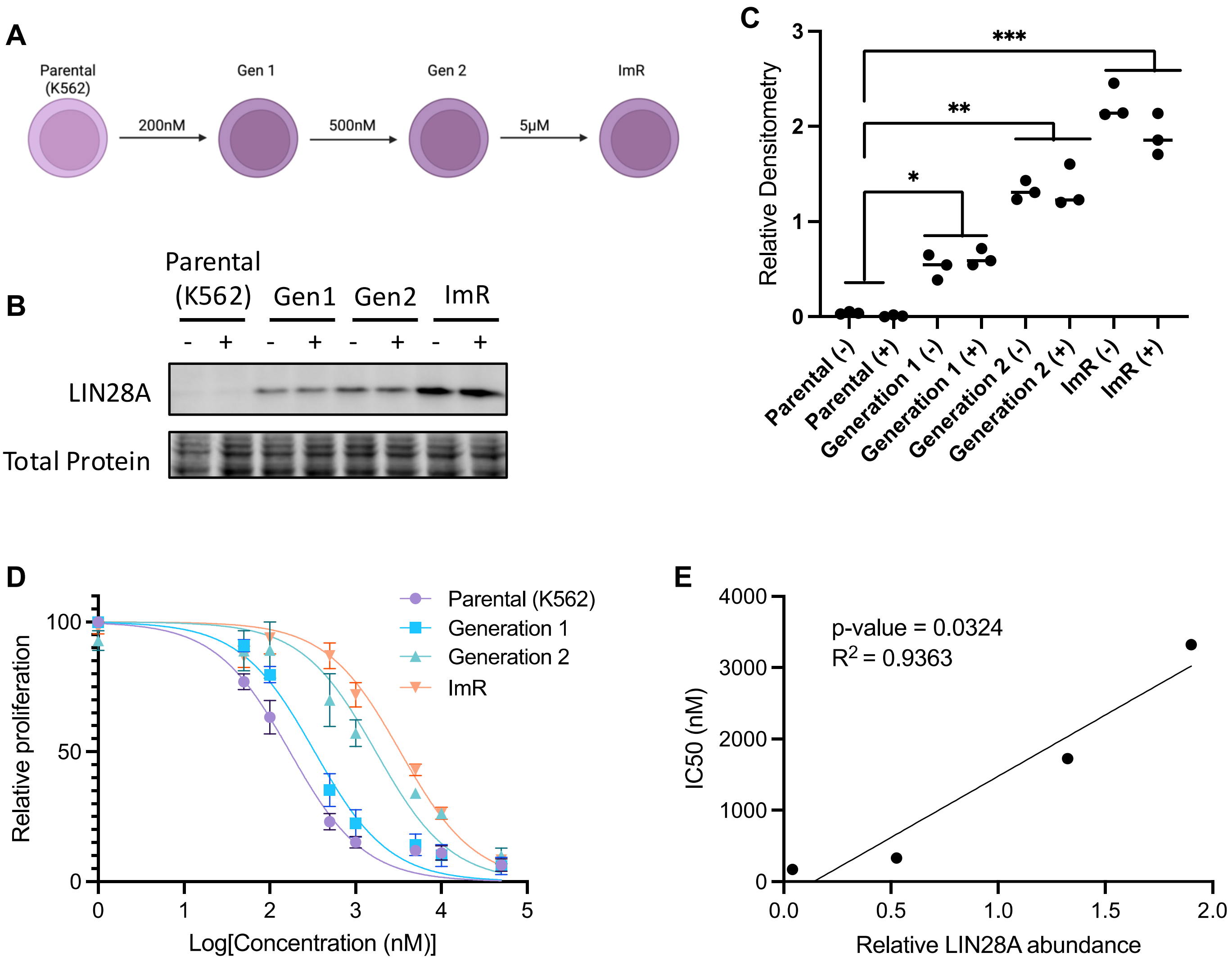


**Figure S8. Progressive development of Imatinib resistance correlates with LIN28A upregulation in K562 cells. (A)** Experimental timeline for generating imatinib-resistant clones through stepwise dose escalation. **(B-C)** Western blot analysis (B) and quantitative densitometry (C) demonstrating gradual LIN28A upregulation across parental K562 cells, intermediate clones (Gen1, Gen2), and the fully resistant ImR line. **(D)** Dose-response curves showing incremental increases in imatinib resistance (Gen1 < Gen2 < ImR). **(E)** Significant positive correlation ($R^2 = 0.9363$) between LIN28A expression levels and imatinib IC50 values across all cell lines. Data represent mean ± SEM (n=3 biological replicates). Statistical significance was determined by one-way ANOVA with Tukey's post-hoc test (*p<0.05, **p<0.01, ***p<0.001).

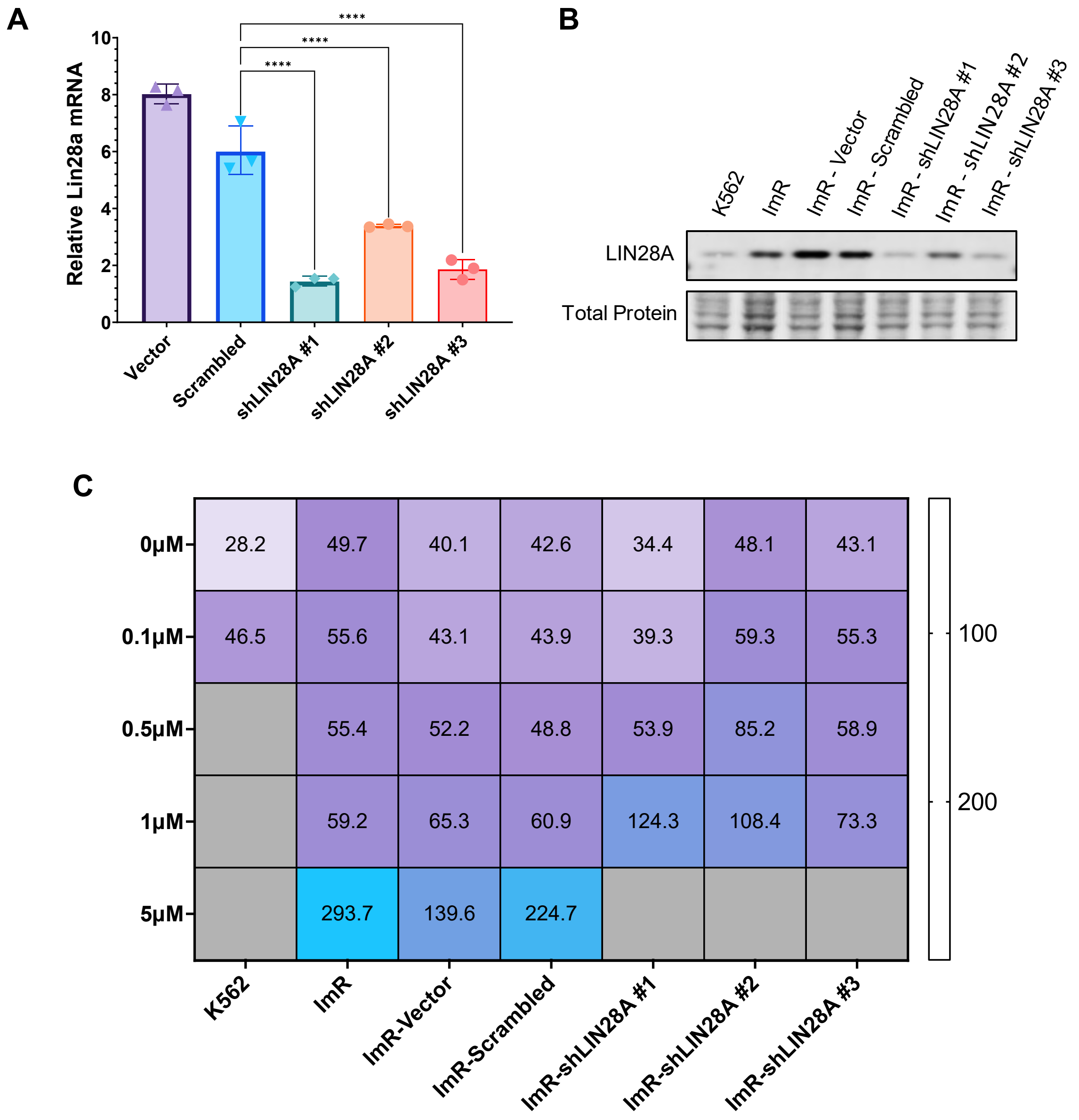


**Figure S9. LIN28A depletion re-sensitizes ImR cells to imatinib treatment.** (**A**) Relative LIN28A mRNA levels normalized against HRPT1 and RLP13A in ImR cells transfected with LIN28A-specific shRNA, scrambled shRNA or vector (mean ± SEM, n=3; ***, $p<0.001$; ANOVA with Tukey post hoc test). (**B**) Western blot confirming LIN28A knockdown in ImR cells using total protein as loading control. (**C**) Doubling time (hr) of the same cells as in (A) when cultured in medium containing increasing concentrations of imatinib.

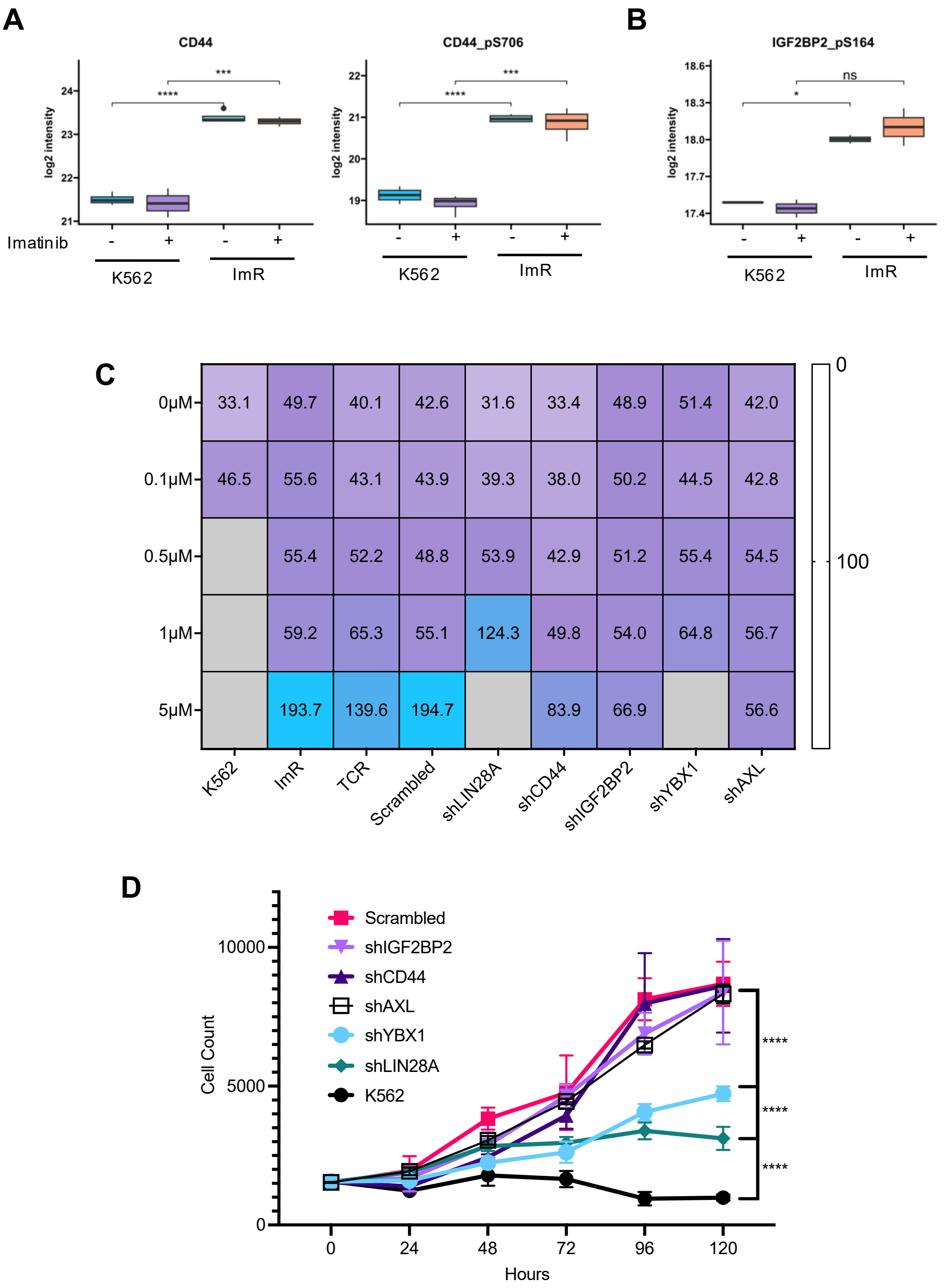


**Figure S10. LIN28A and YBX1 are involved in imatinib resistance.** (**A, B**) Box plots showing significant increases in CD44, CD44-pS706, and IGF2BP2-pS164 in ImR compared to K562 cells. +/-, cells cultured with or without 1 μM imatinib. (**C**) Heatmap showing doubling times of the same cells as in (C) upon treatment with increasing concentrations of imatinib (100 nM to 10 μM). (**D**) Representative growth curves of ImR cells in which LIN28A, CD44, IGF2BP2, YBX1, or AXL was depleted by the corresponding shRNA. Cells were allowed to grow in 1μM Imatinib for 120 hours and the cell numbers were counted every 24 hr. ****, p<0.001; ANOVA with Tukey post hoc test.

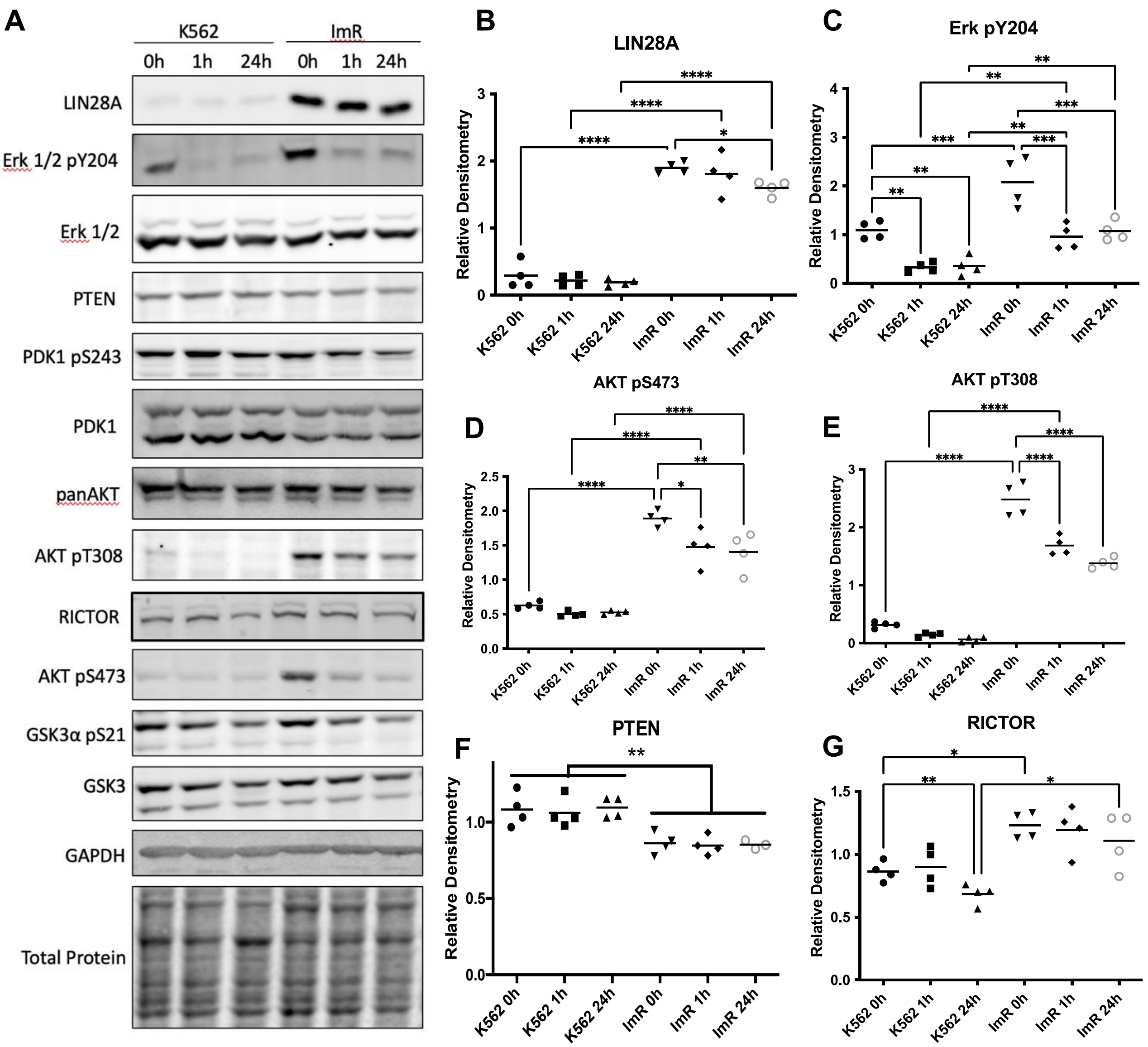


**Figure S11. Increased Lin28A levels were correlated with elevated AKT and ERK signaling in ImR cells.** (**A**) Representative Western blots showing alterations in expression or phosphorylation of LIN28A, Erk1/2, and components of the PI3K-AKT signaling pathway between K562 or ImR cells upon treatment with 1 µM imatinib for various durations (n=4). (**B-G**) Quantification of the Western blot data in (A) for LIN28A (B), Erk1/2-pY204 (C), AKT-pS473 (D), AKT-pT308 (E), PTEN (F) and RICTOR (G). Mean ± SEM; *, p<0.05; **, p<0.01; ***, p<0.001; ***, p<0.001; ANOVA with Tukey post hoc test.

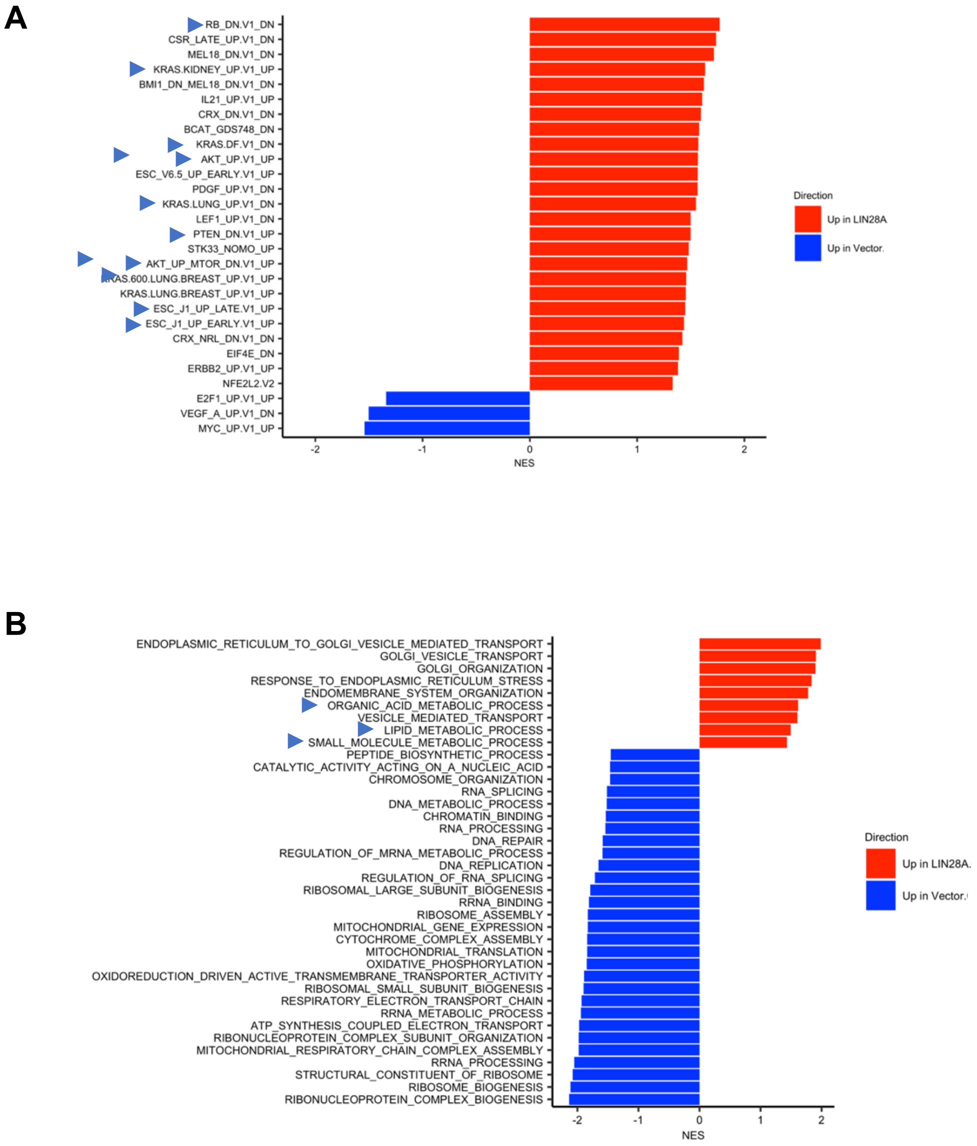


**Figure S12. Overexpression of LIN28A in K562 cell alters oncogenic and metabolic pathways.** GSEA identified significant changes in Oncogenic Signatures (**A**) and Gene Ontology (GO) biological processes (**B**). Red and blue bars denote significant upregulation and downregulation in the LIN28A$^{OE}$ compared to K562 cells. Data shown are based on MS analysis of cells cultured in the absence of imatinib. Blue arrows in (B) denote significant changes that were also identified in ImR cells.

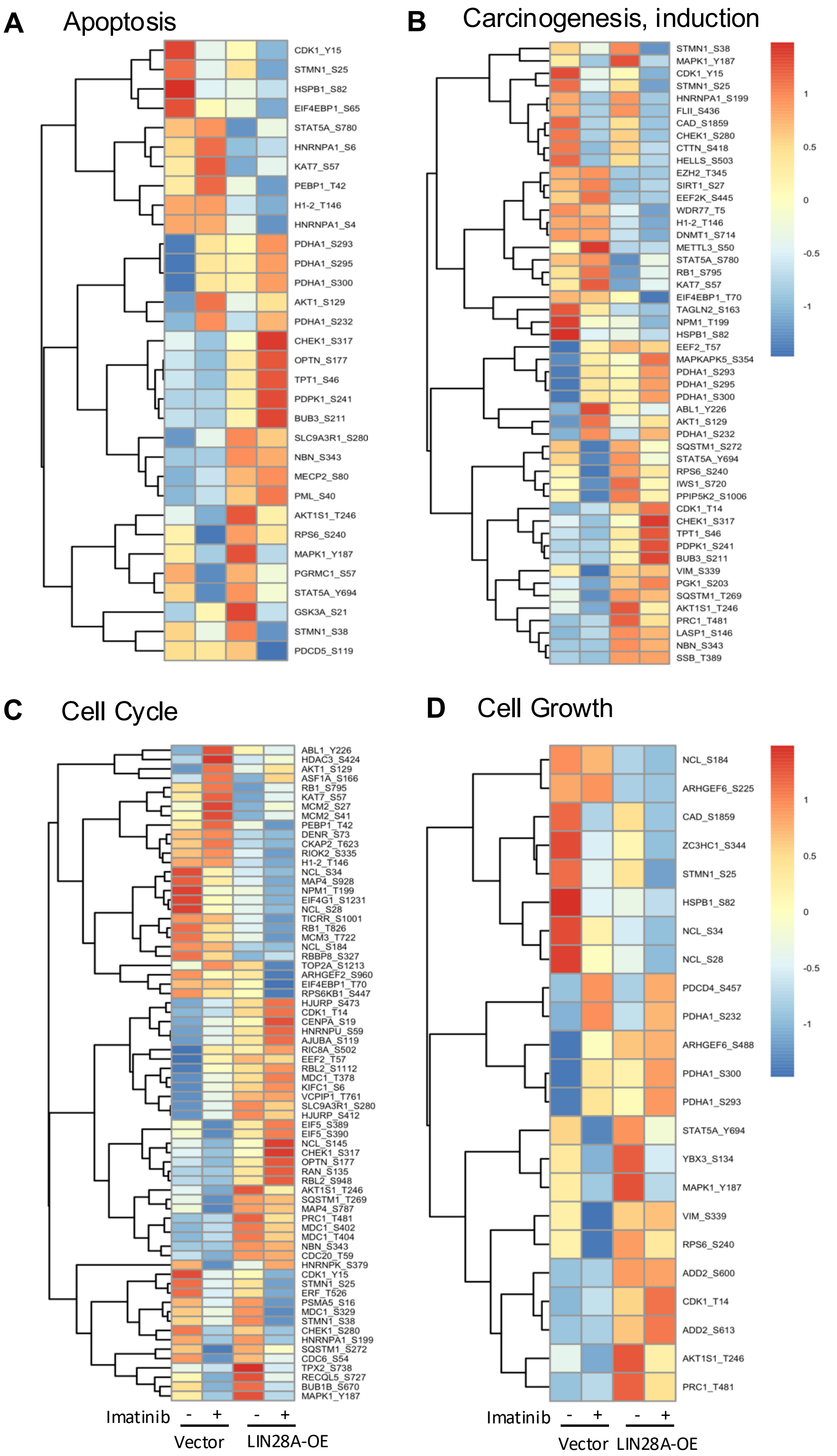


**Figure S13. LIN28A overexpression (OE) alters phosphorylation signalling associated with apoptosis, carcinogenesis, cell cycle, and cell growth.** Significantly changed phosphosites were clustered using Pheatmap based on the corresponding z scores. Proteins shown were from PSP annotation comparing K562 and ImR, with (+) or without (-) 500 nM imatinib.

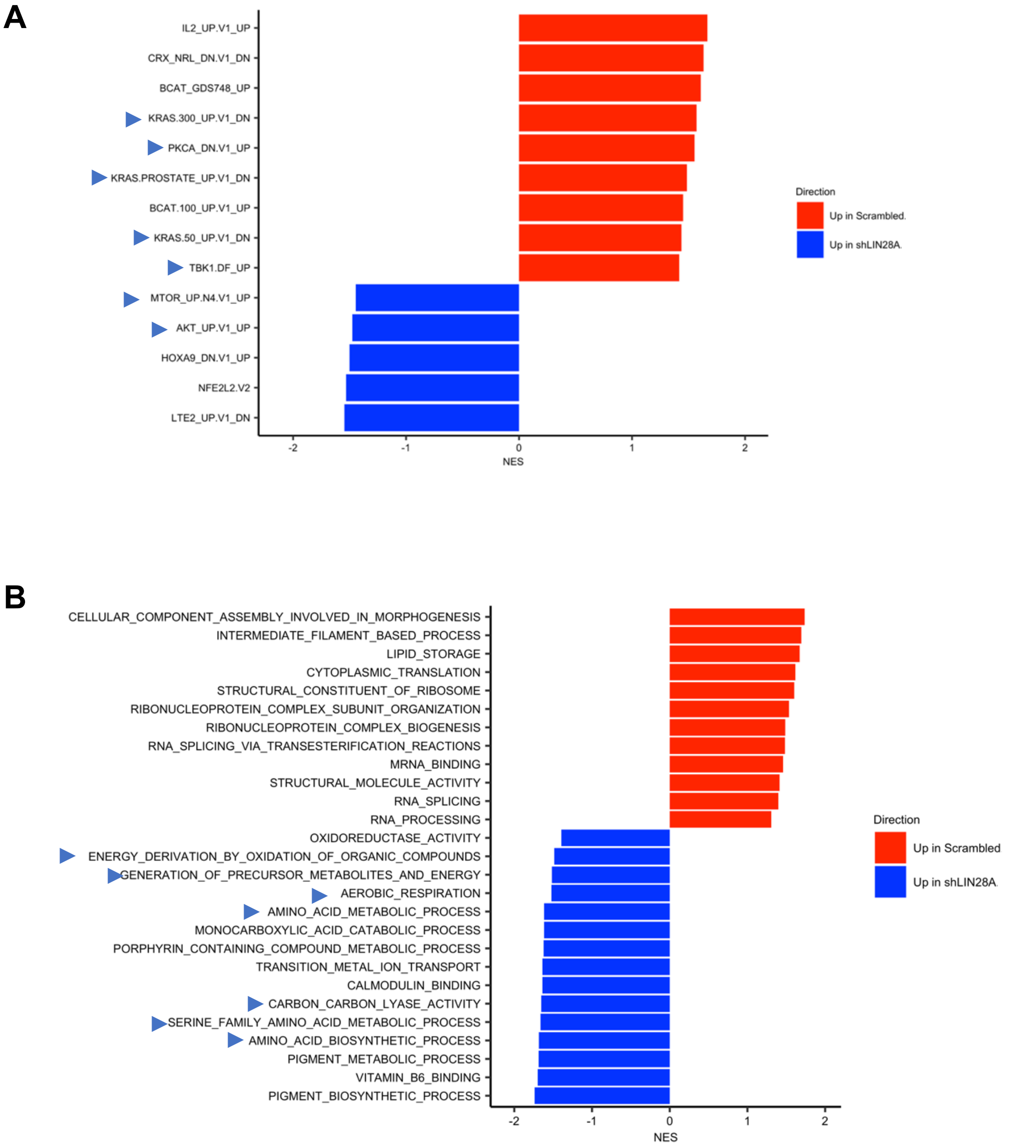


**Figure S14. Knockdown of LIN28A from ImR led to significant changes in oncogenic and metabolic pathways.** GSEA identified significant changes in oncogenic signatures (**A**) and Gene Ontology (GO) biological processes (**B**). Red and blue bars denote significant upregulation and downregulation in ImR cells transfected with the scrambled shRNA vs. LIN28A-specific shRNA. Data shown are based on MS analysis of cells cultured without imatinib. Blue arrows denote oncogenic signatures that were also identified in ImR cells, compared to K562 cells.

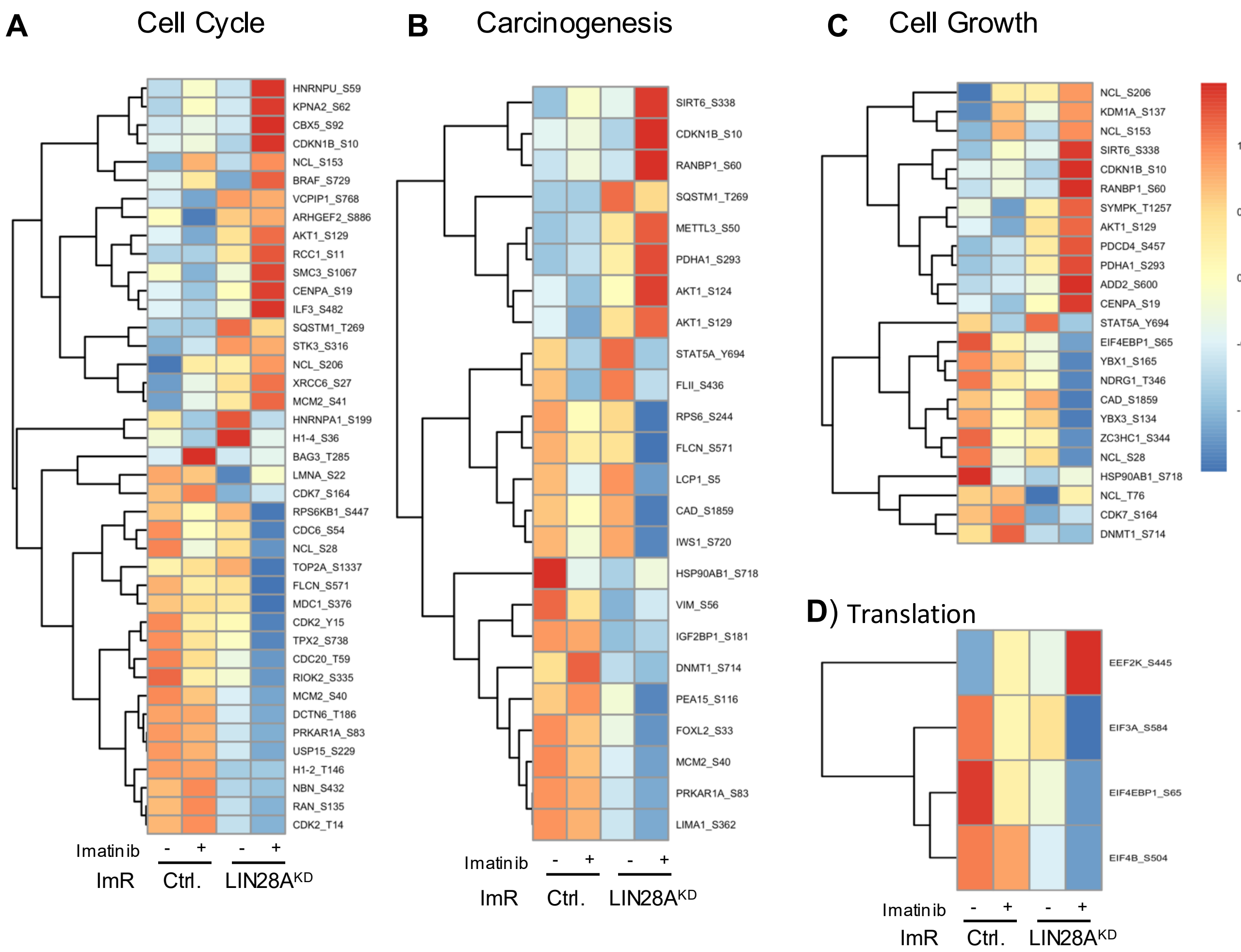


**Figure S15. Alterations in phosphorylation signalling in response to LIN28A knockdown.** Heatmaps showing significantly altered phosphosites (based on z-score) clustered using Pheatmap. Proteins annotated by PSP regulatory sites, comparing ImR cells transfected with LIN28A-specific shRNA (LIN28A$^{KD}$) or a scrambled control (Ctrl) are shown for cell cycle (A), induction of carcinogenesis (B), cell cycle (C) and translation (D). +/- cells cultured in the presence or absence of 500 nM imatinib.

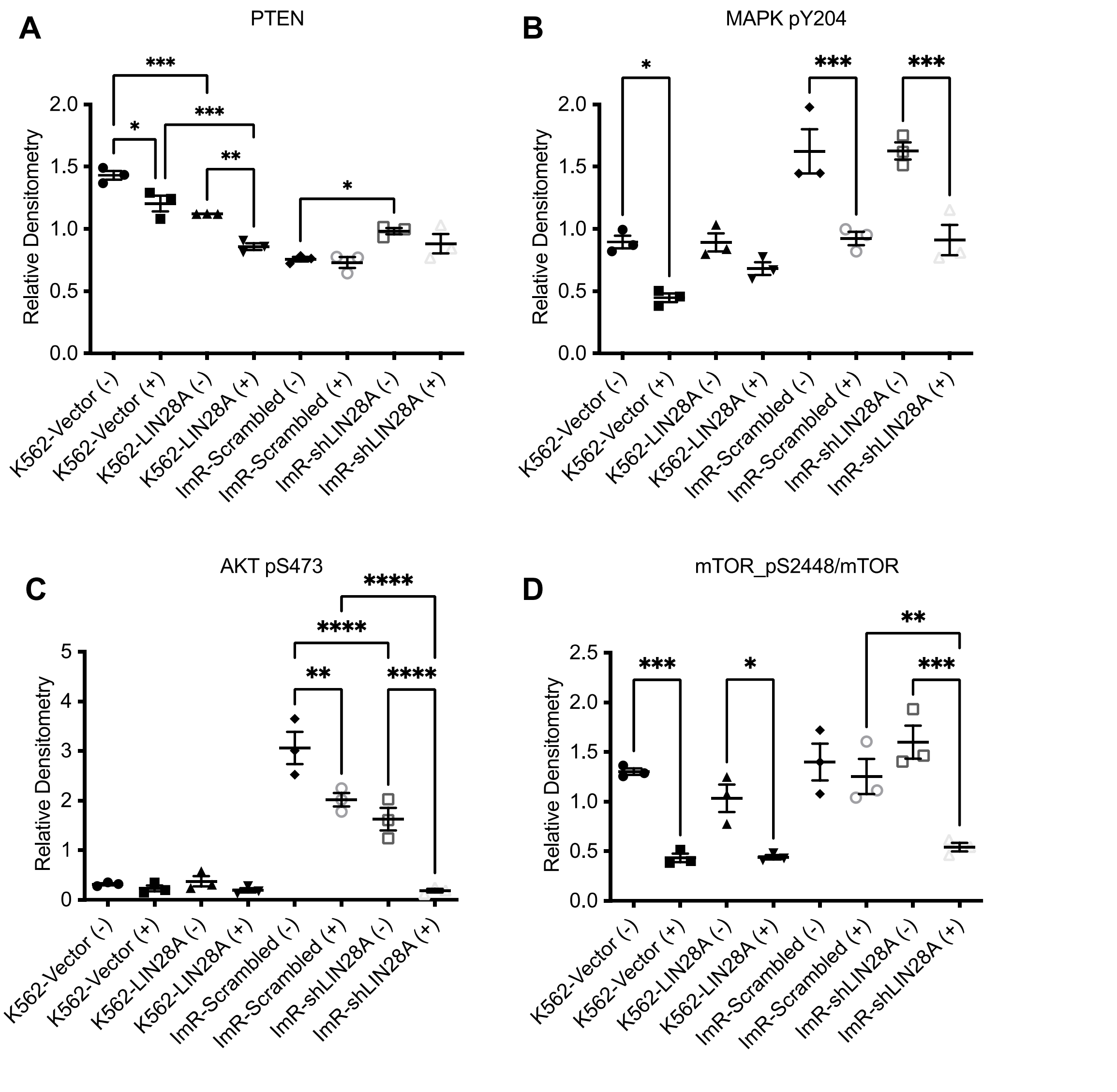


**Figure S16. Alterations in LIN28 expression led to changes in AKT-mTOR signalling.** (A-D) Densitometry quantification of Western blots in Figure 5F showing changes in AKT and mTOR phosphorylation in response to LIN28A overexpression in K562 or knockdown in ImR (cultured +/- imatinib for 24 hrs). *, $p<0.05$; **, $p<0.01$; ***, $p<0.001$; ****, $p<0.0001$; ANOVA with Tukey post hoc test.

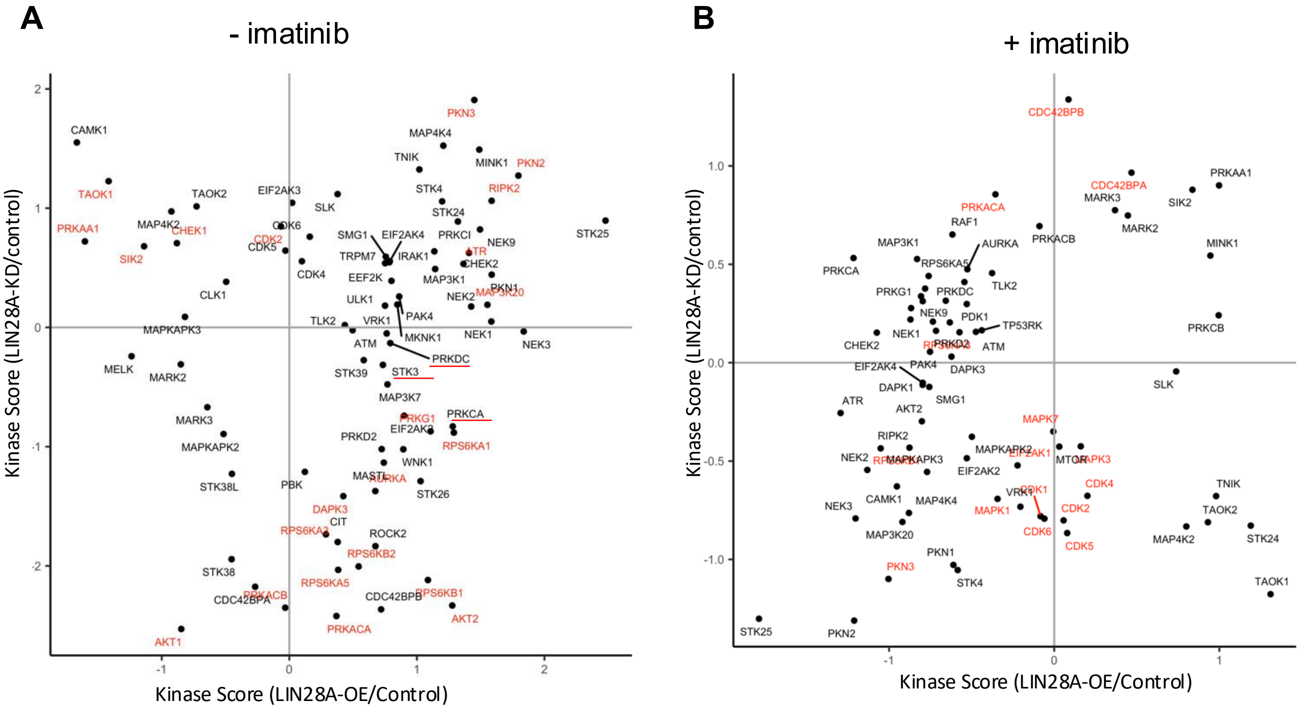


**Figure S17. Cellular LIN28A levels affect STK activity.** Shown are kinase scores predicted by Kinase Library from phosphoproteomic data comparing K562-LIN28A$^{OE}$ with vector control (X axis) and ImR-LIN28A$^{KD}$ with scrambled shRNA control (Y-axis). Kinase highlighted in red also exhibited significant differences between K562 and ImR cells.

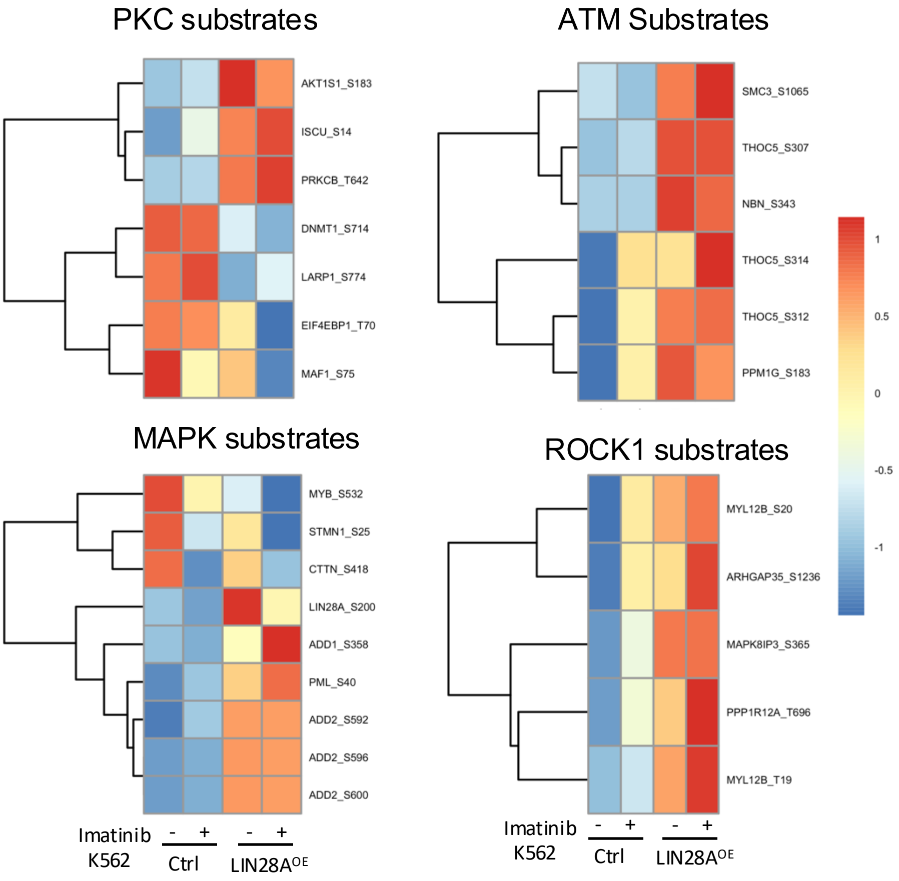


**Figure S18. Alteration in substrate phosphorylation for select Ser/Thr kinases in response to LIN28A overexpression in K562 cells.** Significantly changed phosphosites were clustered using Pheatmap based on the corresponding z scores.

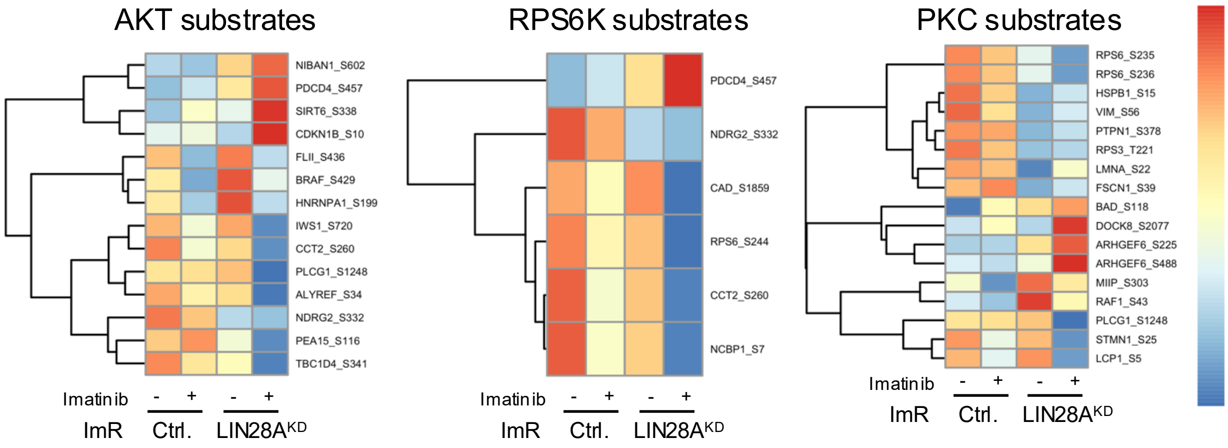


**Figure S19. Alterations in substrate phosphorylation for AKT, PRS6K, and PKC in response to LIN28A knockdown from ImR cells.** Significantly altered phosphosites (based on z-score) were clustered using Pheatmap.

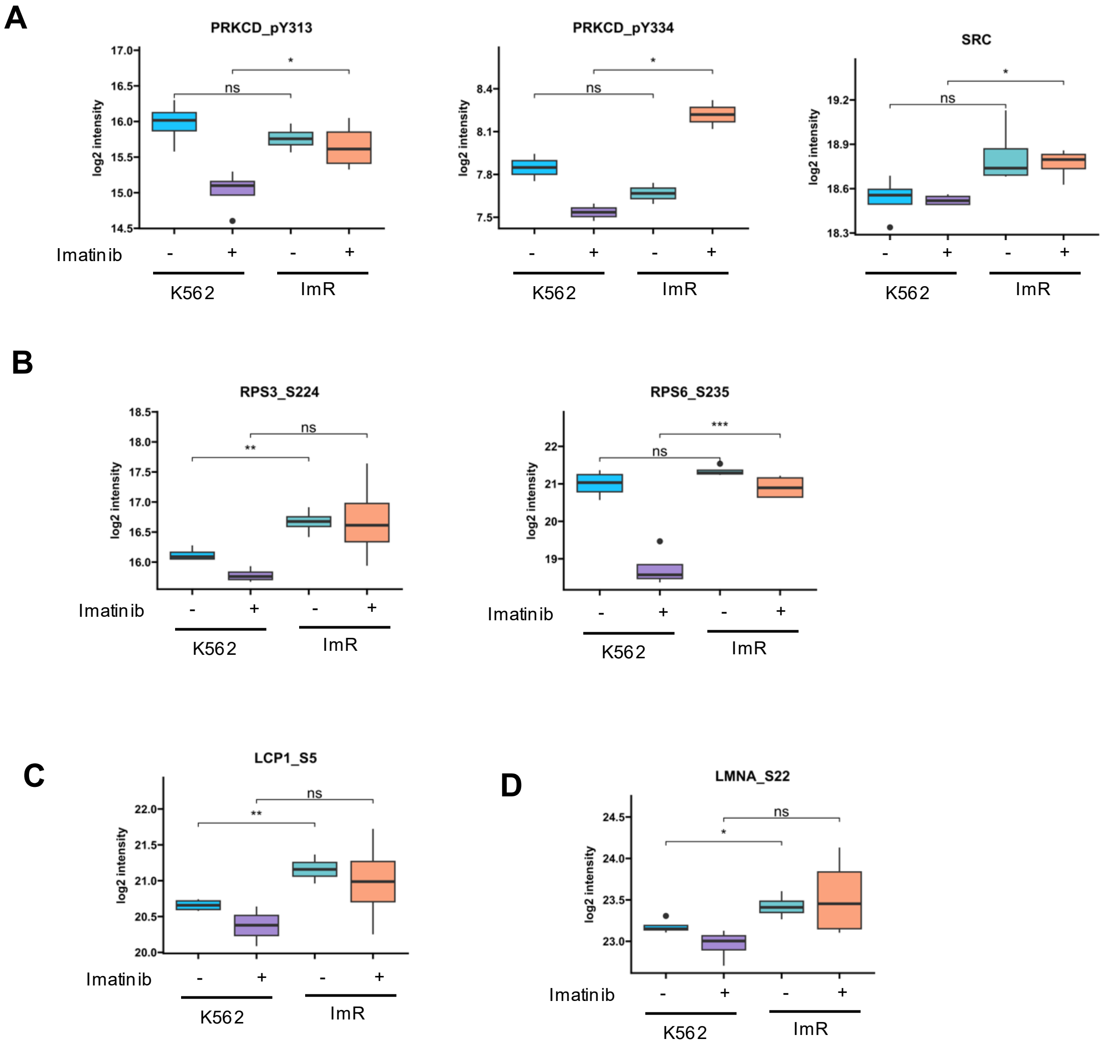


**Figure S20. PKCδ (PRKCD) regulates LCP1, LMNA, and RPS3/6 phosphorylation in an LIN28A-dependent manner.** Compared to K562, phosphorylation of PRKCD (PKCδ) and its upstream kinase Src (A), and substrates RPS3 and RPS6 (B), as well as LCP (C) and LMNA (D), was significantly increased in ImR cells. *, $p<0.05$; Student's t-test.
